# A Shh/Gli-driven three-node timer motif controls temporal identity and fate of neural stem cells

**DOI:** 10.1101/809418

**Authors:** José M. Dias, Zhanna Alekseenko, Ashwini Jeggari, Marcelo Boareto, Jannik Vollmer, Mariya Kozhevnikova, Hui Wang, Michael P. Matise, Andrey Alexeyenko, Dagmar Iber, Johan Ericson

**Affiliations:** Department of Cell and Molecular Biology, Karolinska Institutet, S-171 77 Stockholm, Sweden; D-BSSE, ETH Zürich, Mattenstrasse 26, 4058 Basel, Switzerland; Swiss Institute of Bioinformatics (SIB), Mattenstrasse 26, 4058 Basel, Switzerland; Department of Neuroscience and Cell Biology, Rutgers-Robert Wood Johnson Medical School, 675 Hoes Lane, Piscataway, New Jersey, 08854, USA; Department of Microbiology, Tumor and Cell Biology, Karolinska Institutet, Stockholm, Sweden; National Bioinformatics Infrastructure Sweden, Science for Life Laboratory, Box 1031, 17121, Solna, Sweden

**Author notes:** Correspondence to Johan Ericson. These authors contributed equally to this study (José M. Dias, Zhanna Alekseenko).

## Abstract

How time is measured by neural stem cells during temporal neurogenesis has remained unresolved. By combining experiments and computational modelling, we here define a Shh/Gli-driven three-node timer underlying the sequential generation of motor neurons (MNs) and serotonergic neurons in the brainstem. The timer is founded on temporal decline of Gli-activator and Gli-repressor activities established through downregulation of Gli transcription. The circuitry conforms an incoherent feedforward loop, whereby Gli proteins promote expression of Phox2b and thereby MN-fate, but also account for a delayed activation of a self-promoting Tgfβ-node triggering a fate switch by repressing Phox2b. Hysteresis and spatial averaging by diffusion of Tgfβ counteracts noise and increases temporal accuracy at the population level. Our study defines how time is reliably encoded during the sequential specification of neurons.

## Main text

Time is a central axis of information during embryogenesis but few mechanisms explaining the timing of developmental events have been resolved^1–4^. In the forming central nervous system (CNS), defined pools of multipotent neural stem cells (NSCs) produce distinct cell types in a specific sequential order and over defined timeframes. In this process, ageing NSCs become progressively restricted in their developmental potential by losing competence to generate early-born cell types^5^ and genome-wide analyses have revealed that NSCs undergo dynamic transcriptional changes over time^6^. However, the composition and functional properties of time-encoding circuitries determining timeframes and point of transitions have not been resolved in any model system^7,8^. Temporal neural patterning in vertebrates is a slow process progressing over days or even weeks depending on species^9,10^. Yet, in several lineages, progenitors undergo remarkably coordinated and fast temporal transitions^9–13^. Biological timers regulating temporal neurogenesis are therefore likely to exhibit properties that counterbalance noise in regulatory networks, but how this is achieved at the molecular level remains unknown.

Temporal patterning contributes to the generation of neural cell diversity at all axial levels of the CNS, but only few transcription factors (TFs) and/or signaling molecules regulating temporal fate and potency have been defined in various temporal lineages of the vertebrate CNS^11,12^. To approach the question of time, we focused on a relatively well-defined lineage in the ventral brainstem that sequentially produces motor neurons (MNs), serotonergic neurons (5HTNs) and oligodendrocyte precursors (OPCs)^13,14^. The lineage is induced by Sonic hedgehog (Shh) and defined by the expression of the TF Nkx2.2^13^, and the temporal progression of differentiation is easy to monitor as the NSC-pool remains at a fixed position and does not comprise the specification of proliferative intermediate progenitors as in the developing neocortex^12^. Nkx2.2^+^NSCs show otherwise many common features to cortical progenitors, as young NSCs are multipotent and competent to generate early- and late-born neurons, and ageing NSCs become progressively restricted in their potential over time^12,15,16^. Additionally, the transition from early-to-late phases of neurogenesis is governed by late-acting extrinsic signals whose activation is intrinsically programmed within the lineage ^16–18^. Young Nkx2.2^+^ NSCs co-express early- and late-acting fate determinants^13,16,19–21^, but the activity of the TF Phox2b predominates by specifying MN fate^16,21^. Once Phox2b is downregulated or genetically ablated MN production is terminated and 5HTNs are generated by default^13^ suggesting that Phox2b functions as temporal effector output. Activators of Phox2b have not been defined, but a self-sustained and temporally delayed activation of Tgfβ operates as an extrinsic signal that triggers MN-to-5HTN fate switch by repressing Phox2b^16^. Thus, Phox2b and Tgfβ are important regulatory components of a timer circuitry, but to understand how time is set by the network, it is necessary to define activators of Phox2b and resolve how the temporally gated activation of Tgfβ is mechanistically implemented. Here, we define a Shh/Gli-driven three-node circuitry which explains how time is encoded in the Nkx2.2^+^ lineage, and identify the intrinsically programmed activation of extracellular switch signals as a means to counterbalance noise by spatial averaging.

### Shh/Gli signaling promotes expression of Phox2b

The sequential specification of MNs, 5HTNs and OPCs by Nkx2.2^+^ NSCs is recapitulated in mouse embryonic stem cell (ESC) cultures in response to timed activation of Shh and retinoic acid signaling^16^. To define genome-wide transcriptional changes over time we determined the transcriptome of Nkx2.2^+^ NSCs isolated at different time points by RNA-sequencing. Since the temporal patterning process is slow and progresses over days^16^, cells isolated at a given time were examined at the population level in order to even out changes of gene expression associated with cell cycle progression and neuronal differentiation^22^. We hypothesized that activators driving *Phox2b* are progressively lost over time since Phox2b becomes downregulated and cells undergo a MN-to-5HTN fate switch in the absence of Tgfβ signaling, but on a delayed temporal schedule^16^. Therefore, we determined genes downregulated between 3.5DDC, when Phox2b expression and MN-production has been initiated, and 5.5DDC when Phox2b becomes downregulated in NSCs (Fig.1a)^16^. ∼2200 genes showed a significant decline over this timeframe (p≤0.05; Log_2_(FC)≥0.22). The number of genes was dramatically reduced with increased fold change (FC) stringency and only 27 genes remained at a FC of ∼5.6 (Log_2_(FC)≥2.5) (Extended Data Fig. 1a). Among these were *Gli1, Gli2* and *Gli3* (Extended Data Fig. 1a) which encode zinc-finger TFs that transduce Shh signaling within the nuclei^23^.

**Fig. 1.**
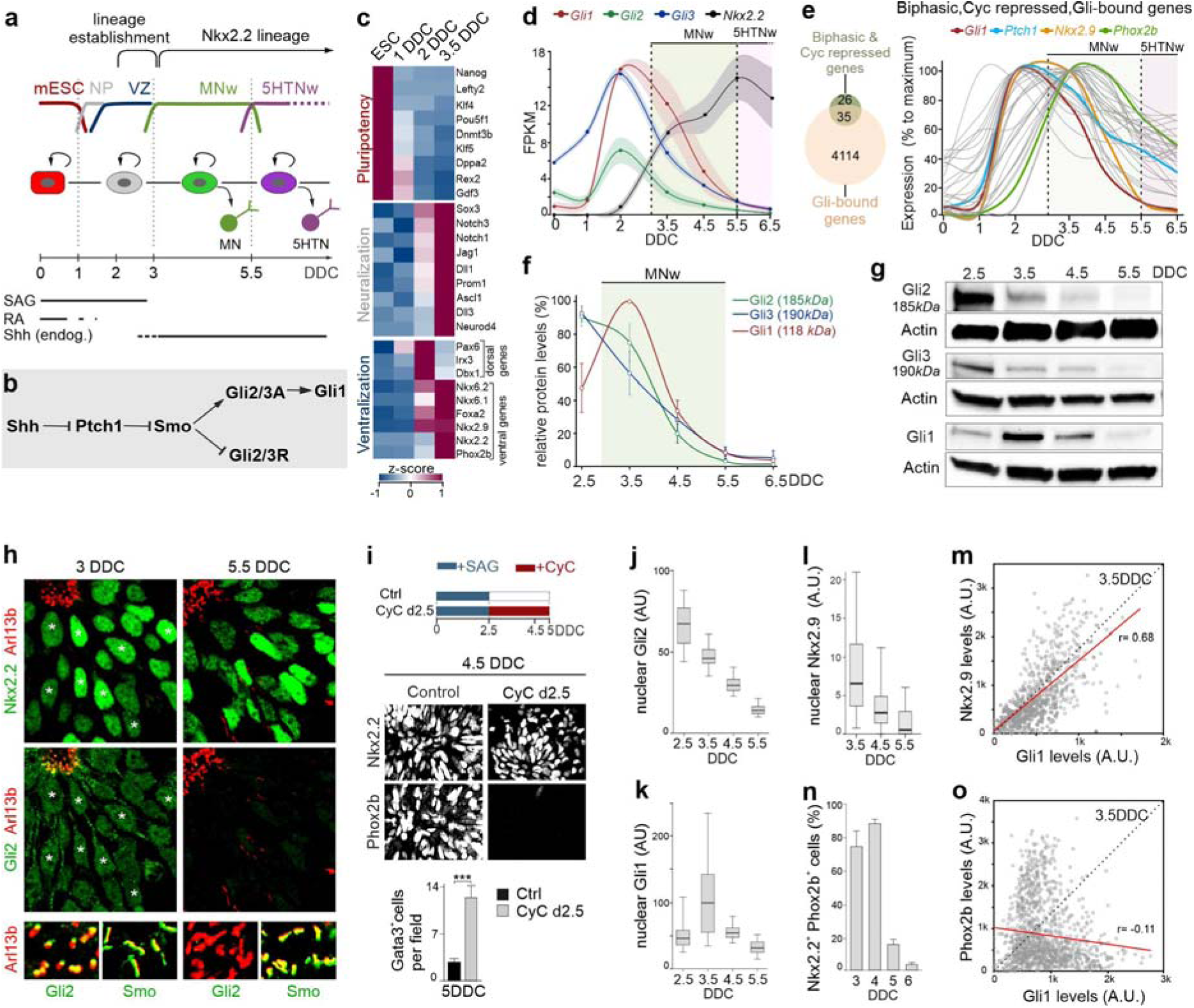
Gli proteins are temporal regulators of Phox2b. **a**, Scheme of mESC differentiation protocol. **b**, Scheme of Shh signaling pathway. **c**, Heat map of expression levels of genes associated with Pluripotency, Neuralization and Ventralization. **d**, Expression levels in FPKM of *Gli1, Gli2, Gli3* and *Nkx2.2* in NSCs over time. Shaded area, S.D. **e**, Genes with biphasic expression profile, repressed at 3.5DDC by Cyclopamine (CyC) treatment (from 0-3.5DDC) and bound by Gli1 or Gli3 proteins, and their relative temporal expression profile. **f,g**, Western blot for Gli1-3 proteins in NSCs at different DDC (**g**) and corresponding quantification (**f**). **h**, Immunofluorescence of Arl13b with Nkx2.2, Gli2 (images correspond to a triple immunofluorescence) and of Arl13b with Smo at 3 and 5.5DDC. **i**, Effect of treatment of cultures with CyC from 2.5DDC on Nkx2.2 and Phox2b expression at 4.5DDC and number of Gata3^+^5HTNs at 5DDC. **j-l**, Box plot of Gli2, Gli1 and Nkx2.9 expression levels in Nkx2.2^+^ nuclei at different DDCs. Whiskers define 5^th^ and 95^th^ percentile, outliers are omitted. **n**, Quantification of Nkx2.2^+^ NSCs expressing Phox2b. **m,o**, Scatter plot of nuclear protein levels of Gli1 with Nkx2.9 (**m**) or Phox2b (**o**) in Nkx2.2^+^ nuclei at 3.5DDC. Trend line in red; r, correlation coefficient. **i,n**, Error bars, mean±SD; Asterisks, Student’s *t* test, *** p≤0.001. MNw, motor-neuron window; 5HTNw, serotonergic-neuron window; DDC, days in differentiation conditions; A.U., arbitrary units.

Gli2 and Gli3 are bifunctional TFs that are processed into repressors (GliR) in the absence of Shh but stabilized as full-length activators (GliA) within cilia by Smo in response to binding of Shh to Ptch1 (Fig. 1b)^23^. Gli1, in turn, is an obligate activator and direct target of the Shh pathway (Fig. 1b) and is together with Ptch1 commonly used as an indicator of ongoing Shh signaling^24^. *Gli1* and *Ptch1* exhibited a biphasic temporal expression: they were upregulated during ventralization of NSCs and induction of the Nkx2.2^+^ lineage (∼1-3DDC) followed by a progressive decline over the Phox2b^+^ MN-window (3-5.5DDC) (Fig. 1c-e). *Gli2* and *Gli3* showed similar biphasic behavior, but were expressed at low levels at ESC-stages (Fig. 1d). We identified 61 genes with biphasic expression similar to *Gli1* and whose induction was inhibited by early treatment of cells with the Smo antagonist Cyclopamine (CyC) (Fig.1e, Extended Data Table 1). Gli1/3 bound^25,26^ in proximity to ∼57% (35) of these genes (Fig.1e, Extended Data Table 1), including *Phox2b*, and *Nkx2.9* which encodes a Shh-regulated TF transiently expressed by Nkx2.2^+^ NSCs^27^ (Fig. 1e). This identifies Gli1-3 as putative activators of Phox2b expression.

Biochemical analyses showed that the downregulation of Gli transcription was translated into a progressive decay of full-length activator forms of Gli2 (Gli2A-185*kDa*) and Gli3 (Gli3A-190*kDa*) between 2.5-5.5DDC (Fig. 1f,g). Gli1 showed a similar profile but reached peak-values at 3.5DDC, consistent with the notion that Gli1 is a target of Gli2/3A (Figs. 1f,g). Gli1-3 proteins were detected in nuclei and Arl13b^+^ cilia at 3DDC but not at 5.5DDC (Fig.1h, Extended Data Fig. 1c,d). Smo translocates to cilia in response to binding of Shh to Ptch1^23^ and localized to Arl13b^+^cilia between 3-5.5DDC (Fig. 1h). Thus, downregulation of Gli transcription mediates a progressive desensitization of the Shh pathway through loss of GliA, despite that Shh/Smo signaling is active over this period. The progressive loss of Gli1-3 expression correlated with the downregulation of Phox2b and Nkx2.9 (Extended Data Fig. 1b). Inhibition of Shh/Smo signaling by CyC subsequent to establishment of the Nkx2.2^+^ lineage resulted in premature downregulation of Phox2b and Nkx2.9 in Nkx2.2^+^ NSCs and increased generation of Gata3^+^ 5HTNs (Fig. 1i, data not shown). These data show that Shh/Gli signaling is required for sustained expression of Phox2b and Nkx2.9 and suggest that inhibition of Shh signaling is sufficient to trigger a MN-to-5HTN fate switch.

Despite the overall decline of Gli proteins over time, there was notable fluctuations of Gli1 and Gli2 expression levels between individual Nkx2.2^+^ nuclei at a given time examined (Fig. 1j,k). Gli1 is a readout of Shh signaling activity^28^ and there was a direct correlation between Nkx2.9 and Gli1 expression levels at 3.5DDC (Fig. 1m). Expression of Nkx2.9 moreover declined over time in a manner similar to Gli proteins (Fig.1j-l, Extended Data Fig. 1b), suggesting a GliA dose-dependent regulation of Nkx2.9. In contrast, there was no correlation between Phox2b and Gli1 expression levels (Fig. 1o), implying that regulators in addition to GliA are likely to influence Phox2b expression. Nevertheless, the fraction of Nkx2.2^+^ NSCs expressing Phox2b remained largely constant between 3-4.5DDC and only dropped dramatically at 5-6DDC when Gli expression approached undetectable levels (Fig. 1j,k,n; Extended Data Fig. 1b) implying that the GliA level sufficient to sustain *Phox2b* transcription is remarkably low.

### Ptch1-independent establishment of parallel temporal GliA and GliR gradients

Immunoprecipitation assays revealed the presence of processed repressor forms of Gli2 (75*kDa*; Gli2R) and Gli3 (83*kDa*; Gli3R) which declined similarly to their respective full-length activator forms between 2.5-5.5DDC (Fig. 2a). Gli3A/Gli3R-ratios remained largely constant while Gli2A/Gli2R-ratios decreased somewhat over time (Fig. 2b). Ptch1 is upregulated in response to Shh signaling which can feedback inhibit the Shh pathway, as free Ptch1 promotes GliR formation through Smo inhibition^23^. Analysis of differentiating Ptch1^-/-^ ESCs showed that the temporal expression profiles of *Gli* genes, Gli1-3 activator proteins, and Nkx2.2, Nkx2.9 and Phox2b were similar in *Ptch1*^**-/-**^ and WT ESC-cultures (Fig. 2c,d, Extended Data Figs. 1b, 2a) establishing that Ptch1 function is largely dispensable in the temporal differentiation process. Interestingly, Gli2R/3R were detected in *Ptch1*^-/-^ cultures and at GliA/GliR-ratios similar to controls (Fig. 2e-f). *Ptch2*^29^ was expressed at extremely low levels in control- and *Ptch1*^*-/-*^ cells (Extended Data Fig. 2b,c). Gli2R/3R were present also in *Ptch1*^**-/-**^ ESC-cultures treated with SAG (Fig. 2e), which activates Smo downstream of Ptch1 and Ptch2^29^. These data reveal that a fraction of bifunctional Gli proteins are processed into GliR-forms in conditions of fully activated Smo, and that the amount of Gli2R/3R and Gli2A/3A produced at a given time is determined by the level of *Gli2/3* transcription (Extended Data Fig. 3). Thus, downregulation of Gli genes produces parallel declining GliA and GliR gradients in the lineage (Fig. 2g).

**Fig. 2.**
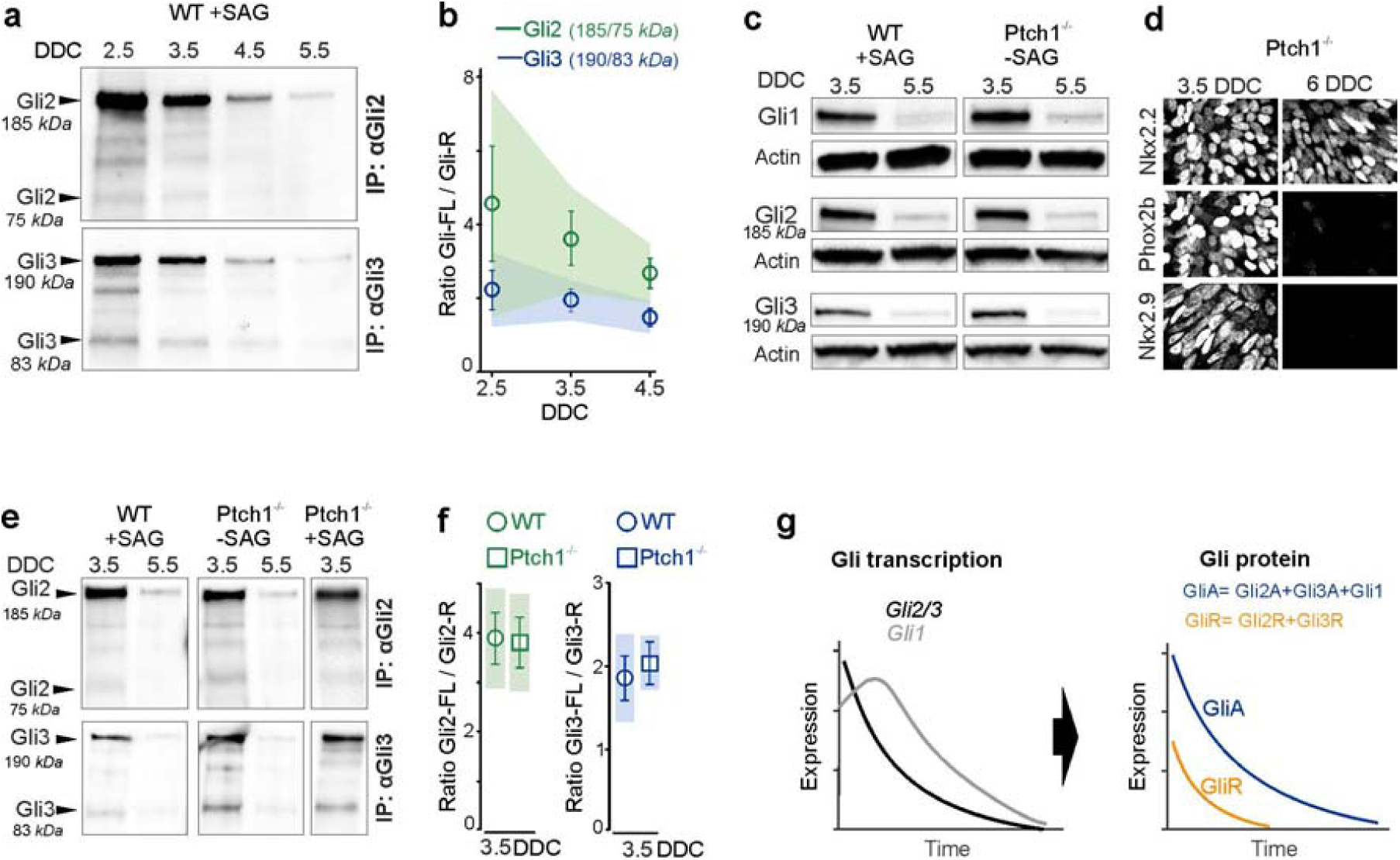
Downregulation of *Gli* genes establishes parallel temporal GliA/GliR gradients. **a,b**, Western blot of immunoprecipitated (IP) Gli2 or Gli3 protein from NSCs isolated at different DDCs and quantification of protein ratios. **c**, Western blot of Gli1-3 in WT and Ptch1^-/-^ NSCs at 3.5 and 5.5DDC. **d**, Immunofluorescence of Nkx2.2, Phox2b and Nkx2.9 in *Ptch1*^-/-^ ESC-cultures at 3.5 and 6DDC. **e,f**, Western blot of immunoprecipitated Gli2 or Gli3 protein from WT or *Ptch1*^-/-^ NSCs isolated at 3.5 and 5.5DDC and differentiated in the presence or absence of SAG as indicated. Quantification of protein ratios of Gli2 and Gli3 bands in WT (+SAG) and *Ptch1*^*-/-*^(-SAG) NSCs at 3.5DDC. **g**, Downregulation of *Gli* genes produces parallel temporal GliA and GliR gradients. **b,f**, Error bars, mean±D; shaded area 95% CIs.

**Fig. 3.**
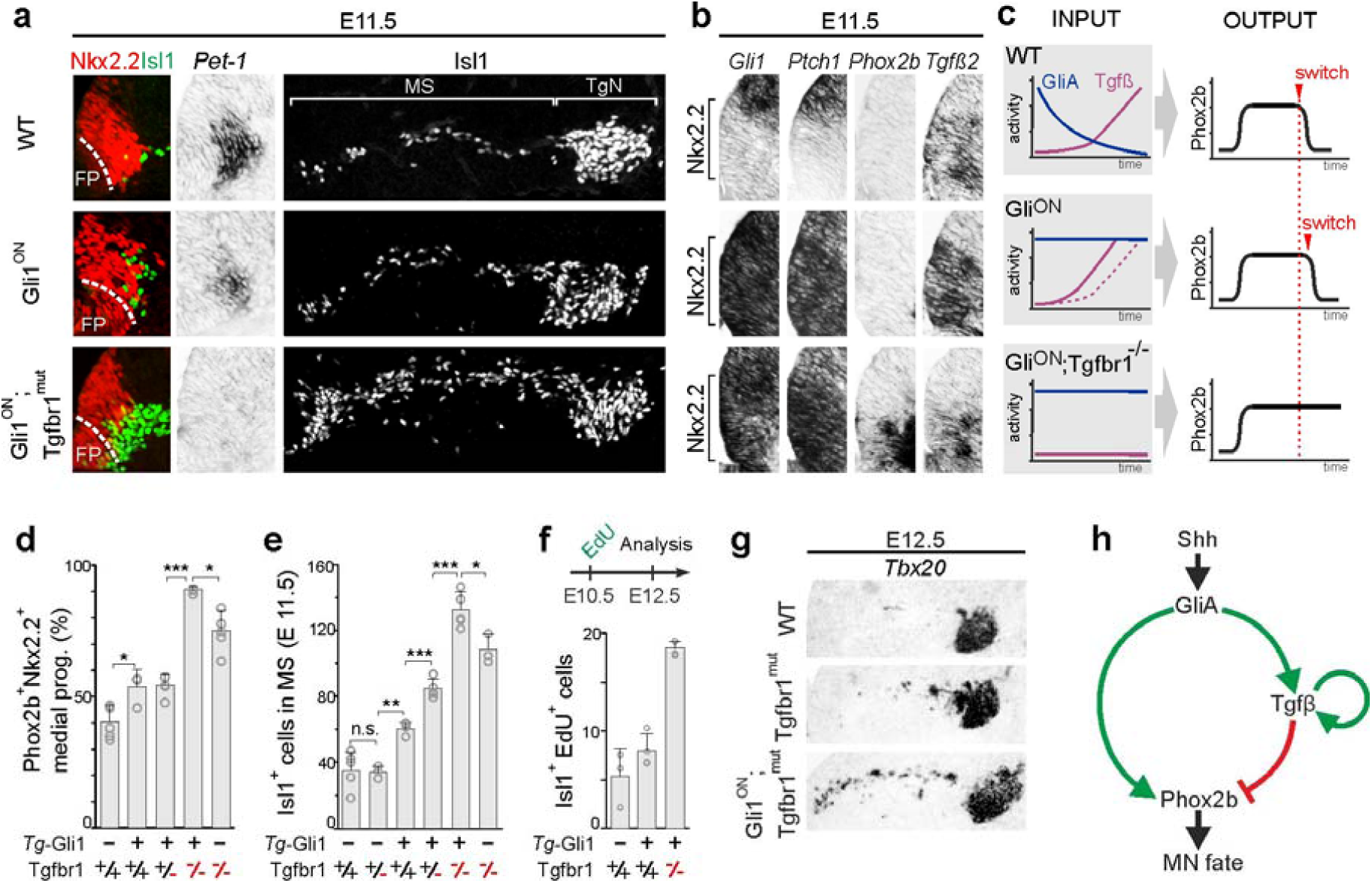
A GliA-driven IFFL network controls the MN-to-5HTN switch time. **a,b,g**, Transverse sections at r2/3 ventral hindbrain level of mouse embryos. **a,b**, Expression of Nkx2.2,Isl1,*Pet-1,Gli1,Ptch1,Phox2b* and *Tgfβ2* at E11.5 in WT and mutant embryos. **c**, Summary of the effects on GliA, Tgfβ and Phox2b expression in different mouse mutants. **d**, Quantification per hemisection of Nkx2.2^+^Phox2b^+^NSCs at E10.5 in WT and mutant backgrounds. Data from wt, *Tgfbr1*^*-/-*^: *n*=5; *Tg-Gli1*^*+*^::*Tgfbr1*^*+/-*^ : *n=*4; *Tg-Gli1*^*+*^::*Tgfbr1*^*+/+*^, *Tg-Gli1*^*+*^::*Tgfbr1*^*-/-*^ : *n=*3 animals per group. **e**, Quantification per hemisection of Isl1^+^-MNs at E11.5 in WT and mutant backgrounds. Data from wt, *Tg-Gli1*^*+*^::*Tgfbr1*^*+/-*^, *Tg-Gli1*^*+*^::*Tgfbr1*^*-/-*^ : *n=*5; *Tg-Gli1*^*+*^::*Tgfbr1*^*+/+*^ : *n=*4; *Tg-Gli1*^*-*^::*Tgfbr1*^*+/-*^, *Tg-Gli1*^*-*^::*Tgfbr1*^*-/-*^ : *n=*3 animals per group. **f**, Quantification of Isl1^+^Edu^+^ cells at E12.5 in WT (*n=*3), Gli1^ON^ (*n=*4) and Gli1^ON^:Tgfbr1^mut^ (*n*=3) embryos pulsed with EdU at E10.5. **g**, Expression of *Tbx20*^+^-MNs at E12.5 in WT, Tgfbr1^mu^ and Gli1^ON^:Tgfbr1^mut^ embryos. **h**, Schematic of the IFFL network. **d-f**, Error bars, mean±SD; Asterisks, Student’s *t* test, * p≤0.05, ** p≤0.01, *** p≤0.001. FP, floorplate; MS, migratory stream; TgN, trigeminal nuclei.

### The temporal network conforms a three-node incoherent feedforward loop circuitry

To define the effect of constant GliA input on temporal output, we generated mice in which Gli1 was constitutively expressed in the Nkx2.2^+^ lineage by crossing a *ROSA26-Gli1*^*FLAG*^ transgene^30^ with a *Nkx6.2-Cre* mouse line^16^ (hereafter termed Gli1^ON^ mice) (Extended Data Fig. 4a). *Ptch1* is downregulated in Nkx2.2^+^ NSCs by E11.5 but was sustained at high levels in Gli1^ON^ mice over this period (Fig. 3b) consistent with continuous GliA expression at high levels. Forced Gli1 expression did not result in any overt spatial patterning phenotype (Fig.3a, Extended Data Fig. 4b) and the number of Isl1^+^ trigeminal MNs was similar to controls at E10.5 (Extended Data Fig. 4a), a time when most MNs have been specified^13^. In controls at E11.5, MN production is terminated and the generation of *Pet1*^+^5HTNs initiated^13^ (Fig. 3a). Most Isl1^+^MNs have migrated laterally to form the trigeminal nuclei but some late-born MNs are still present in the migratory stream (MS) (Fig. 3a), and *Phox2b, Ptch1* and *Gli1* have been downregulated and *Tgfβ2* upregulated in Nkx2.2^+^ NSCs (Fig. 3b). There was a surplus of Isl1^+^ MNs in the MS and reduction of *Pet1*^+^ 5HTNs in Gli1^ON^ mice at E11.5 (Fig. 3a) but the expression of *Phox2b* in Nkx2.2^+^ NSCs was repressed (Fig. 3b). This suggests a mild temporal extension of MN-production in Gli1^ON^ mice, but underpins that *Phox2b* is suppressed and cells undergo a MN-to-5HTN fate-switch on an almost normal temporal schedule in conditions of constant GliA input (Fig. 3c).

**Fig. 4.**
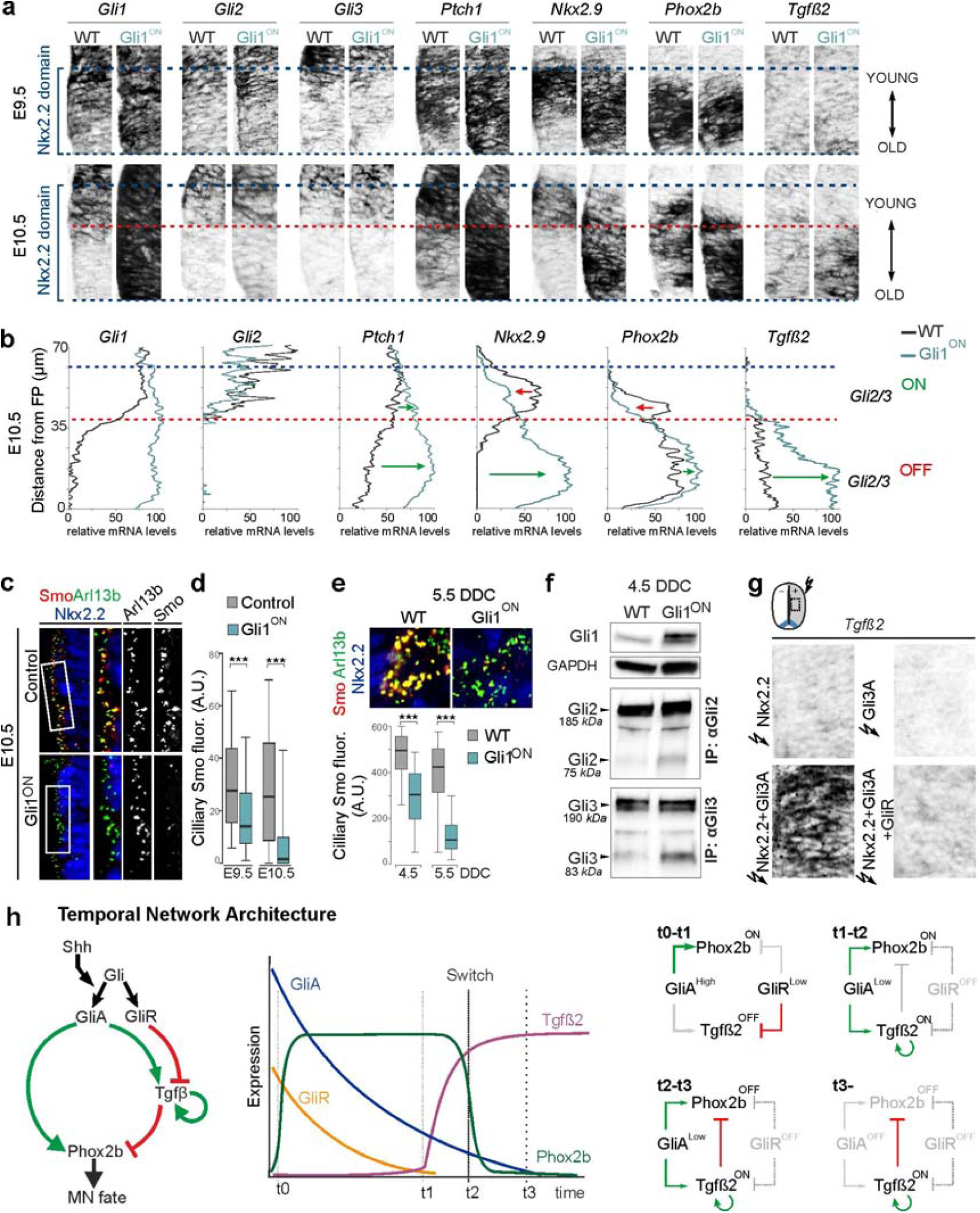
Differential regulation of gene expression by GliA and GliR levels. **a,c**, Transverse sections at r2/3 ventral hindbrain level of mouse embryos. **a,b**, Expression of *Gli1-3,Ptch1,Nkx2.9,Phox2b* and *Tgfβ2* in WT and Gli1^ON^ embryos at E9.5 and E10.5 in Nkx2.2 progenitor domain and plots of relative transcript expression levels along the ventral-dorsal extent of the Nkx2.2 progenitor domain at E10.5. Blue dashed lines delimit Nkx2.2 domain; red dashed line, approximate ventral limit of *Gli2/3* expression. **c,d**, Expression of Smo in Arl13b^+^ cilia within Nkx2.2 domain at E10.5 and boxplots of Smo protein levels in individual Arl13^+^ cilia at E9.5 and E10.5 in WT and Gli1^ON^ embryos. **e**, Expression of Smo, Arl13b and Nkx2.2 in WT and Gli1^ON^ ESC-cultures at 5.5DDC and boxplots of Smo levels in Arl13^+^ cilia at 4.5 and 5.5DDC. **f**, Western blot of Gli1 and of immunoprecipitated Gli2 or Gli3 proteins from WT and Gli^ON^ NSCs isolated at 4.5DDC. **g**, Electroporation of CAGG-constructs expressing Nkx2.2, Gli3A, Nkx2.2+Gli3A, Nkx2.2+Gli3A+GliR at HH12 and analysis of *Tgfβ2* expression in neural tube (boxed region) after 40 hours. **h**, Summary of the temporal network. **d,e**, Boxplots, whiskers define 5^th^ and 95^th^ percentile, outliers are omitted. Asterisks, Pairwise Wilcoxon Test, *** p≤0.001.

Interestingly, the mild temporal phenotype in Gli1^ON^ mice was associated with a marked upregulation of *Tgfβ2* at E11.5 (Fig. 3b), revealing a feedforward propagation of *Tgfβ2* transcription by GliA. This implies a three-node circuitry forming an incoherent feedforward loop (IFFL), whereby GliA activates Phox2b but also the suppressive Tgfβ-node negatively regulating Phox2b (Fig. 3h). This predicts that the MN-window should be extended if GliA is maintained and Tgfβ concurrently inactivated, and to test this we crossed Gli1^ON^ mice onto a *Tgfbr1* mutant background. Removal of one or both copies of *Tgfbr1* resulted in progressively more pronounced temporal phenotypes, as determined by quantification of Phox2b^+^NSCs, Isl1^+^ MNs and by EdU-birth dating experiments (Fig.3d-f, Extended Data Fig. 5a). In *Gli1*^*ON*^:*Tgfbr1*^**-/-**^ at E11.5, there was a massive accumulation of pre-migratory and migrating MNs, a complete lack of *Pet1*^+^ 5HTNs and maintained expression *Phox2b* in NSCs (Fig. 3a,b). The extension of MN production was more pronounced in *Gli1*^*ON*^:*Tgfbr1*^**-/-**^ as compared to *Tgfbr1*^**-/-**^ mutants (Fig. 3e,g, Extended Data Fig. 5b) supporting that the delayed fate switch in *Tgfbr1* mutants^16^ occurs due to depletion of GliA. Additionally, *Tgfβ2* expression was reduced in *Gli1*^*ON*^:*Tgfbr1*^**-/-**^ relative to Gli1^ON^ mice (Fig. 3b) suggesting that positive feedback signaling by Tgfβ is necessary for robust induction of *Tgfβ2* downstream of GliA. Collectively, these data support an IFFL regulatory topology for the GliA-Phox2b-Tgfβ motif (Fig. 3h), and establish that Tgfβ predominates over the Shh pathway by suppressing Phox2b even if cells express GliA at levels sufficient to sustain *Phox2b* transcription.

**Fig. 5.**
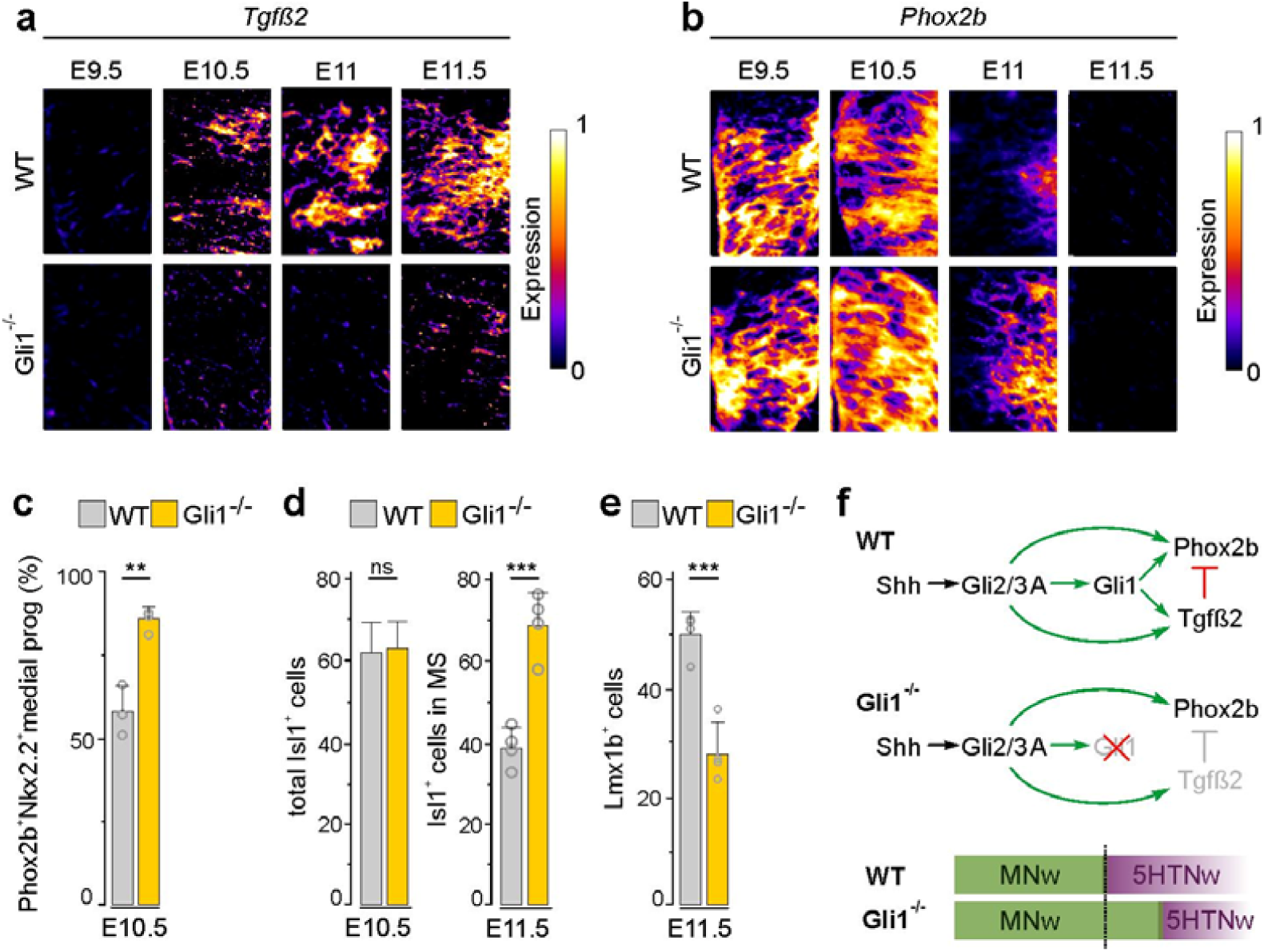
Gli1 is required for *Tgfβ2* induction and correct temporal output. **a,b**, Expression of *Tgfβ2* and *Phox2b* in Nkx2.2 domain at r2/3 level of WT and *Gli1*^-/-^ embryos at indicated stages. **c-e**, Quantification of: medially-located Nkx2.2^+^NSCs expressing Phox2b at E10.5, Isl1^+^MNs at E10.5, Isl1^+^MNs in MS at E11.5 and Lmx1b^+^5HTNs at E11.5 in WT and *Gli1*^-/-^ embryos. Data from wt *n*=3 and *Gli1*^*-/-*^ *n*=4 embryos per genotype at E10.5 and from wt and *Gli1*^-/-^ *n*=4 embryos per genotype at E11.5. Error bars, mean±SD.; Asterisks, Student’s *t* test, ** p≤0.01, *** p≤0.001, ns non-significant. **f**, Summary of response properties of *Tgfβ2* and *Phox2b* to GliA activity.

### Different GliR-sensitivities account for sequential activation of *Phox2b* and *Tgfβ2*

GliA promotes both *Phox2b* and *Tgfβ2* raising the key question how *Tgfβ2* induction can be circumvented at early stages when GliA activity peaks? As the downregulation of *Gli* genes produces parallel declining GliA and GliR temporal gradients, we considered that the delayed activation of *Tgfβ2* could be explained by an inhibitor-titration regulation, a regulatory motif known to convey non-linear switch-like responses^31,32^. To explore this possibility, we characterized the spatiotemporal dynamics of gene expression in Nkx2.2^+^ NSCs in vivo by semi-quantitative in situ hybridization and qPCR. Of note, induction of the Nkx2.2^+^ domain and subsequent MN-to-5HTN fate-switch progress in a ventral-to-dorsal (V→D) manner^13,33,34^ and ventrally located progenitors are thereby older than more dorsally located siblings at a given time. Consistent with this, downregulation of *Gli1-3, Ptch1, Phox2b* and *Nkx2.9* progressed in an overall V→D fashion but at different kinetics (Fig. 4a). *Gli2/3* were most rapidly downregulated and had by E10.5 become constrained to the dorsal third of the Nkx2.2^+^ domain, both in WT and Gli1^ON^ mice (Fig. 4a,b, Extended Data Fig. 6). There was a mutually exclusive relationship between *Tgfβ2* and *Gli*2/3 both in control and Gli1^ON^ mice (Fig. 4b). Upregulation of *Tgfβ2* in Gli1^ON^ mice was first detected at E10.5 in ventral cells that ceased to express *Gli2/3*, and which had initiated *Tgfβ2* expression also in controls but at lower levels (Fig. 4a,b, Extended Data Fig. 6). This shows that a young *Gli2/3*^+^ context is non-permissive for GliA-mediated *Tgfβ2* induction while *Tgfβ2* responds to GliA in a dose-dependent manner in older Gli2/3^-^ cells. Accordingly, *Gli2/3*^+^ cells must express an inhibitor that acts dominant-negative over GliA and that is removed over time, as is the case for GliR (Fig. 2a,g). In functional support for such a repressor function of GliR, we found that forced expression of GliR was sufficient to suppress GliA-mediated activation of *Tgfβ2* in epistasis experiments (Fig. 4g). Removal of GliR is thus necessary for activation of *Tgfβ2* and the kinetics of its elimination therefore determine the kinetics of *Tgfβ2* induction. Due to the bifunctional nature of Gli2 and Gli3 proteins, we have not been successful in selectively eliminating GliR in the Nkx2.2^+^ lineage and cannot therefore rule out that early suppression of *Tgfβ2* may involve repressors in addition to GliR. However, this would not change the overall topology of the timer circuitry, since such hypothetical repressors would then need to be removed either with the same kinetics or faster than GliR to permit activation of *Tgfβ2* when the GliR concentration declines.

**Fig. 6.**
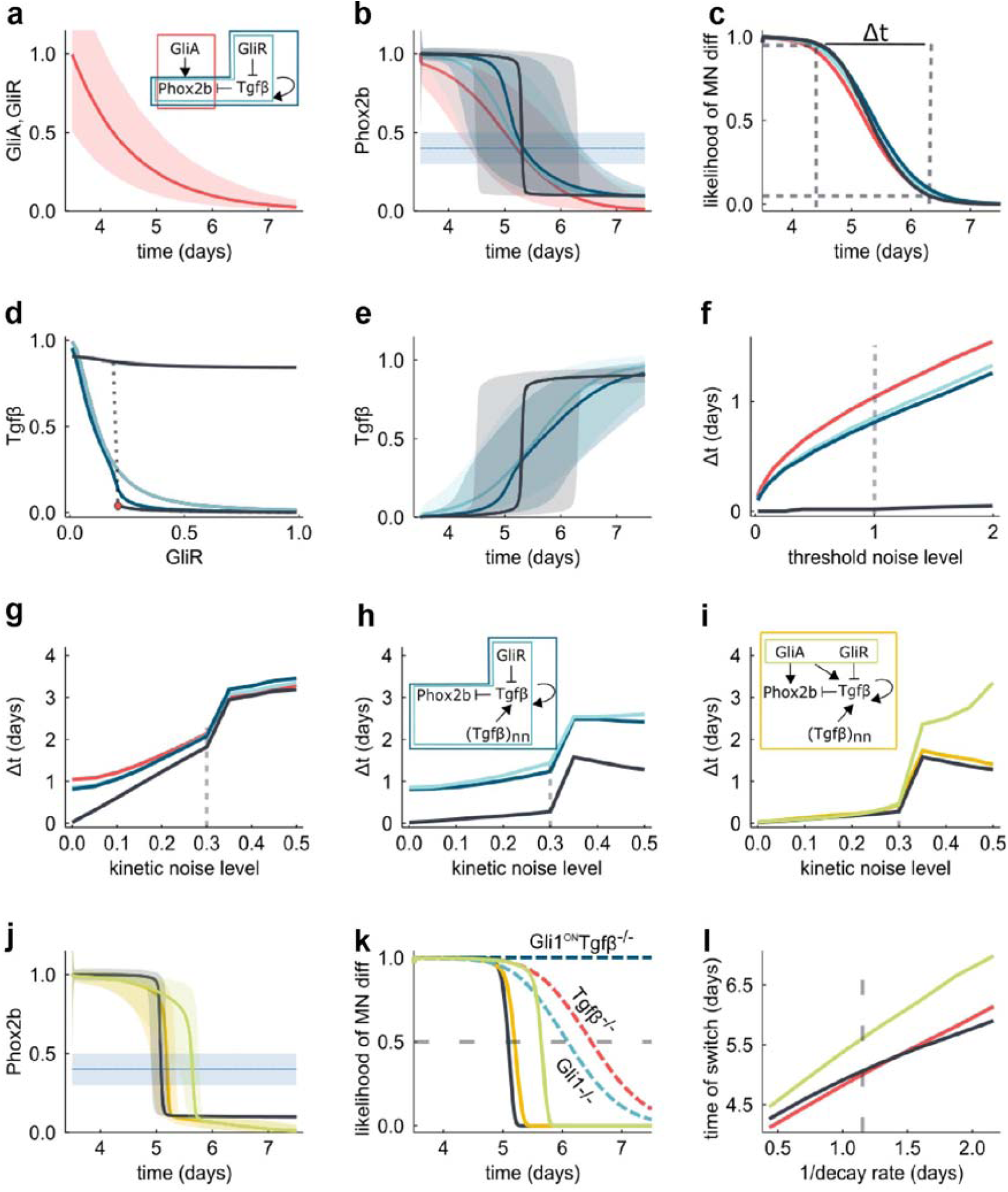
Robust temporal switch by hysteresis and spatial averaging. **a**, GliA and GliR decay exponentially over time. Red line, median values; shaded area, 90% confidence interval for 10^4^ simulations in the presence of kinetic noise. **b**, Predicted concentration of Phox2b over time for different regulatory networks shown in (**a**): Phox2b production is directly activated by GliA (red) or controlled by a GliR-repressive relay (cyan), including Tgf/3 self-activation (blue) and in case of hysteresis (black). Horizontal line and shadow, noisy threshold for motor-neuron (MN) differentiation. **c**, Phox2b-dependent likelihood of MN differentiation over time. Δt, time it takes for the likelihood of MN differentiation to decrease from 0.95 to 0.05. **d**, Bifurcation diagram of Tgf/3 versus GliR showing different sensitivities of models without (light blue) and with (dark blue, black lines) Tgf/3 self-activation. Red dot, bifurcation point for the model with a hysteretic switch (black line). **e**, Predicted concentration of Tgf/3 over time for the different regulatory networks. **f**, Time interval to switch from MN-to-5HTN differentiation (Δt) as a function of the threshold noise level for the different regulatory networks. Vertical dashed line, standard threshold noise used in simulations. **g-i**, Time interval to switch from MN-to-5HTN differentiation (Δt) as a function of the kinetic noise level (**g**) for regulatory networks in panel **a**, (**h**) considering in addition spatial averaging by diffusible Tgf/3, (**i**) and also a positive regulatory link from GliA on Phox2b and Tgf/3 in the hysteretic model (black line), where noise in GliA and GliR is either correlated (green) or uncorrelated (yellow). Vertical dashed lines, standard kinetic noise used in simulations. **j**, Predicted concentration of Phox2b over time for the models in panel **i. k**, Likelihood of MN differentiation over time for the models in panel **i** and indicated mutant mice (dashed lines). **l**, Timepoint of the switch (likelihood of MN differentiation is 0.5) for different GliA and GliR decay rates. Dashed vertical line, standard value used in the simulations.

Unexpectedly, in Gli1^ON^ mice, *Phox2b* and *Nkx2.9* expression was augmented in ventral *Gli2/3*^-^ cells but instead reduced in dorsal *Gli2/3*^+^ cells at E10.5 (Fig. 4b). These transcriptional outputs can only be explained if *Phox2b* and *Nkx2.9* display certain GliR sensitivity, and that the failure to downregulate *Ptch1* in Gli1^ON^ mice results in reinforced Ptch1-mediated feedback inhibition of the Shh pathway. Indeed, we observed a striking displacement of Smo out of cilia in Gli1^ON^ conditions over time (Fig. 4c-e) and increased formation of Gli2/3R in Gli1^ON^ ESC-cultures (Fig. 4f). Thus, increased GliR formation due to Ptch1-mediated feedback begins to suppress *Phox2b* and *Nkx2.9* expression in young Gli2*β*^+^ cells in Gli1^ON^ conditions. This cannot occur once cells ceased to express bifunctional Gli proteins, thereby resulting in the anticipated Gli1-mediated upregulation of Phox2b and Nkx2.9 in older Gli2*β*^-^ cells. Collectively, these data support a model whereby a high GliR-sensitivity for *Tgfβ2* prohibits GliA-mediated activation until GliR has been titrated out, thereby establishing a delayed activation of the Tgfβ-node (Fig. 4h). *Phox2b* and other Shh-regulated genes display lower GliR-sensitivities (*Tgfβ2>Phox2b, Nkx2.9>Ptch1, Gli1*) allowing early activation in the lineage (Extended Data Fig. 7). These data further establish that the decay of GliA followed by downregulation of Gli genes is functionally important, as it is necessary for evading Ptch1-mediated feedback inhibition which otherwise begins to interfere with Phox2b expression during the MN-window.

### Gli1 is required for *Tgfβ2* induction and prompt termination of MN production

A Gli inhibitor-titration regulation of *Tgfβ2* requires that GliA levels remain high enough to activate *Tgfβ2* once GliR has been titrated away, raising the possibility that the feedforward activation of *Gli1* by bifunctional Gli proteins functions to boost GliA activity late in the differentiation process. In direct support for this, we found that *Tgfβ2* failed to be induced in *Gli1*^-/-^ mice (Fig. 5a). This was accompanied by prolonged expression of *Phox2b* in progenitors (Fig. 5b,c), a moderate overproduction of Isl1^+^MNs and concurrent reduction of Lmx1b^+^5HTNs (Fig.5d,e). This shows that Gli1 is required for *Tgfβ2* activation and prompt termination of Phox2b and MN-production (Fig. 5f), suggesting that Gli2/3A are the primary activators of *Phox2b* while induction of *Tgfβ2* instead critically depend on Gli1. Also, considering that Gli1 binds to DNA with lower affinity than Gli2^35^ and *Tgfβ2* is a direct target of Gli proteins^36^, this provides a mechanistic rationale for why *Tgfβ2* is not accessible for Gli1-mediated induction until Gli2/3R are depleted.

### Robust temporal switch by hysteresis and spatial averaging

The timer motif suggested by experimental data relies on the parallel temporal decay of a repressor and an activator, and the delayed activation of a diffusible repressor (Fig.4h). For all progenitors to undergo a coordinated MN-to-5HTN switch in a short time window, cells have to downregulate Phox2b synchronously, despite that Gli protein levels vary between cells (Fig. 1j,k). If we model declining GliA and GliR levels with similar experimental noise levels and a noisy read-out threshold (Fig. 6a,b), we predict a noisy decline of Phox2b (Fig. 6b) and a transition period from MN-to-5HTN production that lasts almost two days (Fig. 6c), both if we consider a direct regulation of Phox2b by GliA (Fig. 6b,c; red line), or a relay via GliR and Tgfβ (Fig. 6b,c; cyan line); we ignore the direct negative impact of GliR on Phox2b as it must be weak, given the strong expression of Phox2b while GliR levels are high. Tgfβ is self-activating, and the introduction of this positive feedback steepens the response (Fig. 6b, blue line), especially when the positive feedback results in a hysteretic switch (Fig. 6b, black line), but does not shorten the transition period (Fig. 6c). In case of a hysteretic switch, the system is bistable, and as the GliR levels decline the system reaches a critical point (Fig. 6d, red dot) where it jumps from a low to a high Tgfβ steady state (Fig. 6d,e; black line). The critical GliR level at which the system returns to the low Tgfβ steady state is at a much higher GliR level, making the switch effectively one-way^37^. Such a hysteretic switch is much more robust to noise in the read-out process (Fig. 6f), but this advantage disappears when also considering the impact of kinetic noise (Fig. 6g) as hysteretic switches are particularly sensitive to kinetic noise (Fig. 6b,e). So, how can the system achieve a fast, reliable transition? Tgfβ is a short range diffusible protein^38^ which permits spatial averaging, and this leads to simultaneous fate switching on the population level despite noise (Fig. 6h). The diffusion range has little impact and even next-neighbor interactions result in robust synchronized switching at the population level. It is important to note that spatial averaging improves the robustness to noise for all network architectures, but the hysteretic switch achieves a substantially shorter transition period (Fig. 6h).

*Phox2b* expression is controlled by both GliA and Tgfβ. Addition of the GliA link makes the switch less robust to noise compared to the GliR link alone (*via* Tgfβ relay), given the lack of spatial averaging. Accordingly, the highest robustness is achieved for a low GliA threshold, as supported by experimental data. Given that generation of Gli2A/3A and Gli2R/3R are coupled (Fig. 2), their levels may be somewhat correlated. However, even perfect correlation (Fig. 6i, green line) does not improve robustness compared to the uncorrelated case (orange), but does lengthen the time to the switch (Fig. 6j,k).

While the timer network is remarkably robust to noise, it remains sensitive to structural changes, which allows us to computationally recapitulate the various mutants discussed above (Fig. 6k), thereby confirming the above verbal reasoning that led to the modelled network architecture (Fig. 4h). The combination of a decaying activator and repressor introduces a further save guard as becomes visible when either component is removed (Fig. 6k). Changes in the Gli decay rate (Fig.6a) alter the time point at which the fate switch occurs (Fig. 6l). In case of an exponential loss of the signal (Fig.6a), the time point changes proportionally to the inverse of the decay rate (Fig.6l).

## Discussion

How time is measured by neural stem cells during temporal patterning of neurons has remained unresolved. In this study, we define a Shh/Gli-driven three-node timer circuitry underlying the sequential specification of MNs and 5HTNs by NSCs. The network is founded on a parallel temporal decay of GliA and GliR established through a progressive downregulation of *Gli1-3* transcription. Regulatory interactions conform an IFFL circuitry in which GliA promotes *Phox2b* expression and MN fate, but also accounts for a delayed activation of a suppressive Tgfβ-node that triggers a MN-to-5HTN fate switch by repressing Phox2b. Data suggest a model in which activation of the Tgfβ-node is temporally gated by a Gli inhibitor-titration mechanism whereby induction of Tgfβ2 by Gli1 is prohibited until GliR has been titrated out. We show that GliR is generated also when the Shh pathway is fully active and downregulation of Gli genes is consequently necessary for elimination of GliR and initiation of a fate switch (Fig. 2,4h). Changes in the rate of Gli decay alter temporal output by the circuitry and conceptually explains how time is encoded in the lineage. In relation to this, Nkx2.2 has been shown to feedback inhibit Gli genes^34,39^ and the MN-window is extended in Nkx2.2^-/-^ mice^13^ suggesting that negative feedback by Nkx2.2 itself impacts on the pace of Gli decay and time-output. Such feedback modulation of Gli genes illustrate how time can be tuned in a lineage-specific manner, which is needed for the generic deployment of the core-timer circuitry in multiple Shh-induced lineages producing distinct neural progenies over different timeframes^16,40,41^.

In the adult brain, the transition of dormant NSCs into transient amplifying cells is accompanied by downregulation of *Gli* genes and is inhibited by GliR^42^ implying that removal of GliR through adaptation of *Gli* transcription accounts for the transition between these cellular states. Together with our study, this suggests that Gli inhibitor-titration and the idea that downregulation of *Gli* genes facilitates induction through removal of GliR may provide a common mechanism to attain switch-like responses in Shh-regulated differentiation processes.

Biological timers based on accumulation or titration of activators or repressors have previously been reported^1,2,4^ and Gli decay is formally sufficient to mediate timer function without Tgfβ, raising the question why timing of the MN-to-5HTN fate switch involves a more complex network architecture. Computational modelling suggest that spatial averaging enabled by the diffusible and self-activating properties of Tgfβ, in combination with hysteresis, produces prompt suppression of *Phox2b* and a coordinated switch at the population level (Fig. 6h,k). Integration of the Tgfβ-node thereby counterbalances noise and generates a more precise timer mechanism as compared to a timer based only on Gli decay. Spatial averaging further provides potential to reset temporal synchrony at the population level which should benefit coordination of subsequent fate transitions as sequentially occurring switches has been shown to be temporally coupled^16^. These community features are not attainable with temporal networks based exclusively on intrinsic transcriptional regulators. Our data consequently provide a functional basis for the intrinsically programmed activation of extrinsic signals in temporal neural patterning processes, and it seems likely that late-acting extrinsic cues implicated in temporal neurogenesis in the cortex^18^ function in a manner analogous to Tgfβ.

## Methods

### Mouse models

The following published mouse strains were used in this study: Nkx6.2::Cre and *Tgfbr1*fl/fl^16^, *ROSA26-Gli1*^*FLAG*^ ^30^, *Gli1*^-/-^ ^43^. Wild-type and experimental mice were maintained in a C57BL/6 background. Male and female mice were both used depending on availability, and were housed in breeding pairs, or group-housed with littermates of the same sex after weaning (2-5 mice/cage), on a 12h light/dark cycle, with food and water provided *ad libitum*. All experimental procedures were conducted in accordance to the Swedish Animal Agency guidelines for animal experimentation and approved by the regional animal ethics committee of Northern Stockholm.

### Mouse ESC derivation

Gli1^ON^ ESCs were derived from E3.5 embryos from crossings between *Nkx6.2::Cre* males and *ROSA26-Gli1*^*FLAG*^ females. Briefly, blastocysts were flushed from uterus and washed twice with KO-DMEM, and plated in ESC derivation medium (KO DMEM, 20% KOSR, non-essential amino acids 0.1 mM, Glutamax 2mM, Penicillin/Streptomycin 100U/ml, ESGRO 1000U/ml and 2-Mercaptoethanol 0.1 mM) containing MEK inhibitor PD0325901 (10µM) on mitotically inactivated mouse embryonic fibroblasts (MEF). In 4-5 days blastocysts attached, inner cell mass outgrowth was dissociated using TrypLE^™^ Express (Life Technologies) and cells were plated in ESC derivation medium on mitotically inactivated MEFs. ESCs were passaged two times on MEFs, following propagation in feeder-free conditions in ESC medium (the same as derivation medium but with 12% KOSR and 3% FCS-ES qualified).

### Differentiation of mESCs into NPCs

Mouse ESCs (E14.1) propagated in ESC media (KO DMEM, 12% KOSR, 3% FCS-ES qualified, non-essential amino acids 0.1mM, Glutamax 2mM, Penicillin/Streptomycin 100U/ml; ESGRO 1000U/ml and 2-Mercaptoethanol 0.1 mM) under feeder-free condition, in a humidified atmosphere containing 5% CO_2_ at 37°C. Stocks of cell lines were free of mycoplasma contamination. For induction of neural differentiation, ESCs were washed twice with PBS, dissociated with TrypLE^™^ Express, passed through a 40 μm cell strainer and seeded at a density of 1-2×10^4^cells/cm^2^ on fibronectin coated surface in N2B27 differentiation medium (50% Neurobasal medium and 50% DMEM/F12, N2 (1:200), B27 (1:100), Glutamax 1mM, Penicillin/Streptomycin 100U/ml, BSA (25µg/ml) and 0.1mM 2-Mercaptoethanol) supplemented with 0.1 μM all-trans RA (Sigma Aldrich) and 0.1 μM Shh-Ag1.3 (SAG). At 3.5DCC, medium was replaced with N2B27 differentiation medium without supplements and changed, subsequently, every second day. To block Shh signaling 2.5 μM Cyclopamine (CyC) was used as mentioned in the text.

### Prominin1-based magnetic sorting (p-MACS) of NPCs

For RNA-seq, qPCR, immunoblotting, and immunoprecipitation experiments NPCs from 3.5DDC and onwards were isolated from differentiating ESCs cultures by Prominin1-based magnetic sorting (p-MACS) according to manufacturer’s instructions (Miltenyi Biotec). Briefly, cells were dissociated using TrypLE^™^ Express, washed twice with ice-cold PBS supplemented with 0.5 % bovine serum albumin and 2mM EDTA (Sigma Aldrich), and passed through a 40 μm cell strainer. Dissociated cells were incubated with magnetic beads conjugated with anti-prominin-1 antibody. Prominin1^+^ cells were separated on magnetic columns and processed for biochemical analysis. To assess purity of sorted fraction, cells were plated, incubated for 2 hours, and fixed for immunocytochemistry.

### Chick electroporation

Fertilized chick eggs were stored up to one week at 16°C, and at the start of experiment eggs were placed horizontally in a 38°C humidified chamber for ∼40 hours until the developing embryo reached a Hammilton-Hamburger stage 11-12. Approximately 5ml of white were removed to lower the embryo and an opening in the shell on the top of the egg was made to expose the embryo which was visualized under a magnifying microscope. A mixture of DNA plasmid(s) and Fasta green (used for coloring) was injected into the central canal of the chick embryo using a pulled capillary needle attached to a mouth aspirator tube. Several drops of PBS were placed on top of the embryos, the electrodes were positioned manually one at each side of the embryos at the hindbrain level and two 25ms, 6V electric pulses 1second apart were applied to the embryo using an Electro square Porator ECM 830. The eggs were sealed and incubated at 38°C for 40 hours before embryo collection and processing for immunofluorescence or *in situ* hybridization.

### Immunohistochemistry

Embryonic stage of collected mouse embryos was determined by somite number (E9.5-E11.5 embryos) and by limb anatomy (E10-E12.5 embryos). Mouse embryos from E9.5-E11.5 5 and chick embryos collected 40 hours post-electroporation were fixed in 4% paraformaldehyde in 0.1M phosphate buffer (PBS) for 2 hour on ice with shaking, and E12.5 mouse embryos were fixed overnight at 4°C. Fixed tissue was washed with PBS, cryoprotected by equilibration in 30% sucrose in PBS, embedded in OCT, frozen on dry ice, and cryostat-sectioned in the transverse plane at 12 μm. Tissue sections were collected at hindbrain level, and rhombomeric level was determined by anatomical landmarks (i.e. otic vesicle) and Phox2b/Isl1 immunohistochemistry to visualize MNs. Immuhistochemistry was carried out in water-humidified chamber. The tissue on slides was incubated with blocking solution (3% FCS/0.1% Triton-X100 in PBS) at room temperature (RT) for 1 hour followed by incubation with primary antibodies overnight at 4°C and fluorophore-conjugated secondary antibodies for 1 hour at RT. Both primary and fluorophore-conjugated secondary antibodies were diluted in blocking solution. Sections were mounted using Vectashield and coverslipped for imaging. For ESC-differentiated material, cells were fixed for 12mins at RT in 4% paraformaldehyde in PBS, rinsed 3 times in PBST (PBS with 0.1% Triton-X100), and blocked for 1 hour at RT with blocking solution (3% FCS/0.1% Triton-X100 in PBS). Cells were then incubated with primary antibodies overnight at 4°C followed by incubation with fluorophore-conjugated secondary antibodies for 1 hour at RT. Primary antibodies used: mouse anti-Nkx2.2, Isl1 (DSHB), mouse anti-Smo, Gata3 (Santa Cruz Biotechnology), mouse anti-Gli1 (Cell signaling Technology), rabbit anti-Arl13b (Proteintech Group), mouse anti-Arl13b (NeuroMab), goat anti-Gli2, Gli3, and Sox1 (R&D Systems), guinea-pig anti-Gli2 (Gift from Dr. Jonathan Eggenschwiler, University of Georgia), guinea-pig anti-Phox2b, Lmx1b, and Nkx2.9 (home made), mouse anti-Actin (Seven Hills Bioreagents), rabbit anti-Arx (Gift from K. Miyabayashi). Sections were mounted using Mowiol/DABCO mounting solution. Images were captured using Zeiss LSM880 confocal microscope.

### *In situ* hybridization

Mounted tissue sections were post-fixed in 4% paraformaldehyde (PFA) in phosphate buffer for 10 minutes (min), washed three times for 5 min with phosphate buffer solution (PBS), incubated with Proteinase K solution (1µg/ml Proteinase K in 50mM TRIS.CL pH7.5, 5mM EDTA) for 5 min and PFA-fixed and washed with PBS as before. Slides were subsequently treated with an acetylating solution (2mM HCl water solution containing 14µl/ml triethanolamide, 2.5µl/ml acetic anhydride) for 10 min, washed three times with PBS and incubated with hybridization solution (50% formamide, 5x SSC, 5x denharts, 250µg/ml Yeast RNA, 500µg/ml Herring sperm DNA, 20mg/ml blocking reagent from Roche) for 1 hour in a chamber humidified with a 50% formamide/5X SSC solution. All steps were performed at RT. Slides were subsequently incubated overnight at 70°C with labelled probe diluted in hybridization solution, followed by washes with a 0.2X SSC solution, first at 70°C for 1h and then by another of 10 min at RT. Slides were incubated in B1 solution (0.1M Tris-Cl pH7.5, 0.15M NaCl, 10% heat inactivated fetal bovine serum) for 1 hour at RT, followed by an overnight incubation with anti-DIG-alkaline phosphatase Fab fragments antibody diluted in B1 solution at 4°C in a water humidified chamber. Excess antibody was washed away with three washes at RT with B1 solution and equilibrated in B3 solution (0.1M Tris-Cl pH 9.5, 0.1M NaCl, 50mM MgCl_2_) before incubation of tissue with developing solution (10% polyvinyl alcohol, 100mM Tris-Cl pH9.5, 100mM NaCl, 5mM MgCl_2_, 0.24 mg/ml levamisol) containing NBT/BCIP (Roche) at RT. The developing reaction was stopped by placing the slides in water and after several washes, slides were mounted using Aquatex mounting medium (Merck). Labelled probes were produced from *in vitro* transcription of linearized plasmid containing a sequence specific for the desired gene using the Dioxigenin (DIG) RNA labeling kit (Roche) according to manufacturer’s protocol. Probes were purified using Microspin G-50 columns (GE Healthcare) and stored at −20°C.

### Edu Labelling

EdU (Life Technologies) was injected intraperitoneally in pregnant female mice at 0.04mg/g of body weight, and detected using Click-iT EdU Alexa Fluor 555 Imaging Kit (Life Technologies) according to manufacturer’s protocol.

### Isolation of mouse neural tissue for qPCR

Neural tube tissue was isolated as described before ^44^. In brief, mouse embryonic tissue was dissected in L-15 medium, staged based on somite number and limb morphology, and hindbrain region was roughly dissected out with the help of tungsten needles. To completely remove the mesenchymal tissue surrounding the neural tissue, the dissected hindbrain piece was treated sequentially with dispase solution (1mg/ml dispase in L-15 medium) for 5 minutes, 10%FBS solution (in L-15 medium) for 5 minutes and washed twice in L-15 medium. The remaining mesenchymal tissue was then removed with the help of tungsten needles and the ventral region of rhombomere 2-3 of the hindbrain was isolated. At the stages analyzed the rhombomeric units of the hindbrain can be easily defined by grooves in the neural tissue. RNA for each individual piece was isolated using RNeasy Micro Kit with a DNAse digestion step (Qiagen) according to manufacturer’s protocol.

### Immunoblotting and immunoprecipitation

For whole cell extracts prominin1-MACS isolated NPCs were lysed in RIPA buffer (Sigma) complemented with protease and phosphatase inhibitor cocktail (Thermo Scientific) and incubated on ice with shaking for 30 min. Lysate was cleared by centrifugation (20 000g for 20 min at 4 °C) and protein concentration determined by Bicinchoninic Acid (BCA) assay. Protein lysate was resuspended in LDS buffer with 2.5% 2-Mercaptoethanol and denaturated at 95 °C for 5 min. 15-30μg of protein were loaded per lane of a 10% SDS polyacrylamide gel (Bio-Rad) and transferred onto nitrocellulose membranes (BioRad) using a Trans-Blot Turbo System (BioRad). Membranes were incubated 1h in blocking solution (TBS with 0.1% Tween-20 (TBST) and 5% nonfat dry milk), followed by overnight incubation at 4 °C with primary antibodies. After 3 washes with TBST, membranes were incubated with HRP-conjugated secondary antibodies for 1h at RT. Detection of HRP was performed by chemiluminescent substrate SuperSignal West Dura substrate and signal was detected on a ChemiDoc Imaging System (Bio-Rad).

For immunoprecipitation assays, 300 μg of protein lysate was diluted with Pierce IP lysis buffer (Thermo Scientific) up to 1ml and incubated overnight at 4 °C with 2µg of primary antibody. Protein-antibody complex was isolated with protein A/G magnetic beads according manufacturer’s instructions (Thermo Fisher Scientific). Precipitated material was used for immunoblotting as described above. Primary antibodies used: mouse anti-Gli1 (Cell signaling Technology), goat anti-Gli2, Gli3 (R&D Systems), mouse anti-GAPDH (Thermo Fisher Scientific) and mouse anti-Actin (Seven Hills Bioreagents).

### Image analysis

For quantification of protein level from immunofluorescence, confocal images were analyzed in Image J 1.48v. Nuclear area was defined by DAPI, Nkx2.2 or Sox1 immunostaining and integrated density of fluorescent signal in individual nuclei was determined.

Images from in situ hybridization were acquired in a Zeiss-AxioImager.M2 microscope coupled to an Axio camera 503 mono and processed in Photoshop CS6. Images in Figures 5A and 5B, were processed in ImageJ 1.48v using LUT-Fire function.

To determine relative gene expression levels along the Nkx2.2 domain (Figure 4), images were analysed in image J 1.48v using “Plot Profile” function. A region of tissue that did not expressed the gene was used to determine background levels.

Western blots were quantified in Image J 1.48v using Gel Analysis Tool.

### RNA extraction, cDNA preparation and qPCR

Total RNA was isolated from cells using RNeasy Mini kit (Qiagen) or Quick-RNA Mini Prep Plus kit (Zymo Research). 500ng of RNA were used for cDNA preparation using Maxima First Strand cDNA synthesis kit (ThermoScientific). Quantitative Real-Time PCR was performed in a 7500 Fast Real Time PCR system thermal cycler with Fast SYBR Green PCR Master Mix (Applied Biosystems). Analysis of gene expression was performed using the 2-ΔΔCt method and relative gene expression was normalized to *Gapdh* transcript levels using primer for mouse: *Gli1* (Fwd: 5’-gtcggaagtc ctattcacgc-3’ and Rev:5’-cagtctgctctcttccctgc-3’), *Gli2* (Fwd:5’-agctccacacacccgcaaca-3’ and Rev: 5’-tgcagctggctcagcatcgt-3’), *Gli3* (Fwd:5’-caaccacagcccttgctttgc-3’ and Rev: 5’-ggcccacccg agctatagttg-3’), *Gli1* (5’UTR) (Fwd:5’-cctttcttgaggttgggatgaag-3’ and Rev: 5’-gcgtctcagggaaggatga-3’), *Ptch1* (Fwd:5’-actgtccagctaccccaatg-3’ and Rev: 5’-catcatgccaaagagctcaa-3’), *Ptch2* (Fwd:5’-cctagaacagctctgggtagaagt-3’ and Rev: 5’-cccagcttctccttggtgta-3’), *Nkx2.9* (Fwd:5’-gtgcgttccacagactg ct-3’ and Rev: 5’-gagtctgcagggcttgtctc-3’), *Phox2b* (Fwd:5’-tgagacgcactaccctgaca-3’ and Rev: 5’-cggttctggaaccacacct-3’), *Gapdh* (Fwd:5’-gtggtgaagcaggcatctga-3’ and Rev: 5’-gccatgtaggccatgaggtc-3’) and *Tgfβ2* (pre-designed from IDT towards exon4/5).

### RNA sequencing, gene expression quantification and differential expression analysis

RNA concentration and integrity were determined on an Agilent RNA 6000 Pico chip, using Agilent 2100 BioAnalyzer (Agilent Technologies). RNA sequencing of condition-specific samples was performed by the National Genomics Infrastructure at the Science For Life Laboratory in Stockholm on Illumina HiSeq2500 in RapidHighOutput mode with single-end setup (1×50 bp read length). Reads were mapped to the mouse genome assembly, build GRCm38 using Tophat (v 2.0.4) ^45^. Next, the number of reads mapped to each gene was calculated using htseq-count (v 0.6.1) ^46^. The gene level abundances were estimated as FPKMs using Cufflinks (v 2.1.1) ^47^. Plots and histograms were made using FPKM values. Further, we processed the read count data with RNA-Seq specific functions of R package limma ^48^. For adjusting the mean-variance dependence we used the function voom ^49^, which rendered the expression data into normally distributed log2-counts per million (logCPM) values, accompanied with observation-level gene-specific adjusting weights. The differential expression was estimated with functions lmFit, eBayes, and topTable. The variance estimates were obtained by treating all samples as replicates (design=NULL) and obtaining library sizes from counts (lib.size=NULL) without further normalization (normalize.method=“none”). The output fold-change values of differential expression were accompanied with p-values. The latter were adjusted for multiple testing by calculating Benjamini-Hochberg’s false discovery rate, FDR ^50^.

### Identification of genes changing over time

To determine genes downregulated in NSCs between 3.5DDC and 5.5DDC, FPKM values at each time point, p-values and fold change (FC) values calculated as described above were used and the following criteria were applied: FPKM (3.5DDC) ≥2, p≤ 0.05 and downregulated (log_2_FC ≥0.22). Additionally, only protein-coding genes were considered.

### Determination of biphasic genes

To identify genes that exhibited a biphasic expression profile during differentiation (from 0-6.5 DDC) and that were repressed by Cyclopamine treatment (CyC) we determined genes whose expression peaked at 2 or 3.5 DDC and were repressed in 3.5DDC progenitors differentiated in CyC conditions. An initial criteria was applied as a cut off of gene expression and Cyc sensitivity (FPKM at 2DDC or 3.5DDC ≥2, and p-value at 3.5DDC between SAG and CyC treated cultures ≤ 0.05, and Log_2_FC at 3.5DDC between SAG and CyC treated cultures ≥0.5 (downregulated)). Next genes whose expression peaked at 2 DDC or 3.5 DCC were determined using the following criteria. Peak at 2DDD: differentially expressed between 2DDC and 6.5DDC and 0DDC (p≤ 0.05), upregulated between 0-2DDC (Log_2_FC(2-0DDC) ≥0.5, Log_2_FC(2-1DDC) ≥0), and downregulated between 2-6.5DDC (Log_2_FC(2-6.5DDC) ≥0.5, Log_2_FC(2-5.5DDC) ≥0.5, Log_2_FC(2-4.5DDC) ≥0). Peak at 3.5DDD: differentially expressed between 3.5DDC and 6.5DDC and 0DDC (p≤ 0.05), upregulated between 0-3.5DDC (Log_2_FC(3.5-0DDC) ≥0.5, Log_2_FC(3.5-1DDC) ≥0, Log_2_FC(2-1DDC) ≥0), and downregulated between 3.5-6.5DDC (Log_2_FC(3.5-6.5DDC) ≥0.5, Log_2_FC(3.5-5.5DDC) ≥0.5). Additionally, only protein-coding genes were considered. The list of genes can be found in Extended Data Table 1.

### Theoretical framework

Our theoretical framework considers the dynamics of GliA (A), GliR (R), Tgf*β* (T) and Phox2b (P). We approximate the measured dynamics of GliA and GliR (Fi. 1N), by an exponential decay from initial values A(0)=R(0) =1:

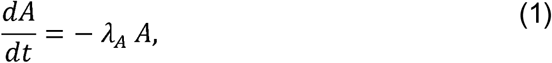

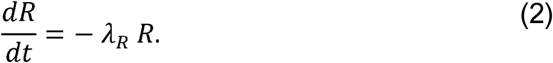

The equations for Tgf*β* and Phox2b differ for the different regulatory networks. A generalized form is given by:

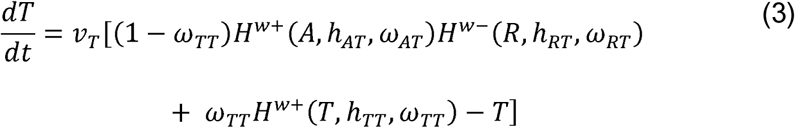

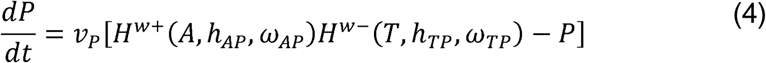

where the functions *H*^*W*+^(*x, h, ω*) = (1-*ω*) + *ω H*^+^(*x, h*) and *H*^*W*−^(*x, h, ω*) = (1−*ω*) + *ω H*^-^(*x, h*) represents weighted positive and negative Hill functions, respectively. For all Hill functions, we use a Hill factor equal to 2, i.e. 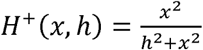 and 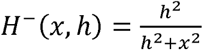. The different links in the regulatory networks are represented via *ω* (Table 1). Thus, in the case of *ω*=0, *H*^*W*+^ = 1 and *H*^*W*−^ = 1, and the link represented by this function is not being considered. In the case of *ω*=1, *H*^*W*+^ = *H*^+^ and *H*^*W*−^ = *H*^−^ represent a positive and negative Hill function, respectively. We use *v*_*T*_ as production and decay rates, so that the maximal concentrations Tgf*β* and Phox2b is one and their values are restricted to the interval [0,1]. To represent the effects of molecular noise, we added scalar white noise to the equations, i.e., a noise term that is proportional to the values of the variables. Therefore, the ordinary differential equations of the form

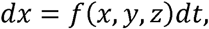

become stochastic differential equations of the form

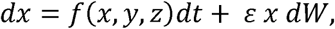

where *W* denotes a Wiener process and *ε* denotes the kinetic noise level.

In Figure 6I-6K, we have considered both correlated and uncorrelated kinetic noise in GliA and GliR. In the correlated case (green lines), we used the same values for GliA and GliR, while in the uncorrelated case (yellow lines), the values are obtained by independently solving Eqs 1,2 with a noise term.

In order to evaluate the impact of noise in the Phox2b concentration on the likelihood of a cell to differentiate into either a MN or 5HTN, we defined the likelihood of MN differentiation as the cumulative distribution function of a gamma distribution with different levels of variance. More specifically, we defined the shape (k) and the scale (*θ*) parameters of the gamma distribution as k = 10/*η* and *θ* = 0.04 *η*, where *η* represents the threshold noise level. The average of this gamma distribution is given by *kθ* = 0.4 for all *η*, while the variance is proportional to *η* and given by 1.6 10^−2^ *η*.

The effect of Tgf*β* diffusion (Figure 6H-K) was introduced by replacing the variable T in the term *H*^*W*+^(*T, h* _*TT*_, *ω* _*TT*_) in Eq. 3 and the term *H*^*W*−^(*T, h* _*TP*_, *ω* _*TP*_) in Eq. 4 with 0.5*T* + 0.5 ⟨ *T* ⟩ _*nn*_, where ⟨ *T* ⟩ _*nn*_ represents the average values of Tgf*β* in the nearest neighboring cells. Averaging over a wider range of neighbours yields similar results. In our simulations, we have considered an array of 100×100 cells with periodic boundary conditions.

#### Mutants

In the table, the full network represents the WT case. For the *Tgfbr1*^-/-^ mutant, we removed the Tgf*β* modulation on Phox2b by setting *ω* _*TP*_ = 0.0. Based on data in Figure 2, we used a 30% lower starting value for GliA in the Gli1^-/-^ mutant compared to WT. Lastly, in the Gli1^ON^Tgfbr1^-/-^ mutant the GliA concentration remains always well above the threshold for Phox2b production. Given that Tgf*β* is absent, the Phox2b concentration always remains above the threshold for MN formation.

#### Parameter values

In the simulations, we have used *λ* _*A*_ = *λ* _*R*_ = 0.86 to approximate the measured decline in GliA and GliR, as well as *v*_*T*_ = 50, and *v*_*P*_ = 90, which set the timescale of the Tgf*β* and Phox2b kinetics, respectively. The other parameter values differ between the models and are presented in the table below. We emphasize that the threshold values *h*_*l*_ were set to reproduce the experimentally observed time point of the switch, but otherwise do not affect the conclusions from the model. Accordingly, we do not present a sensitivity analysis.

Parameter values used in the simulations

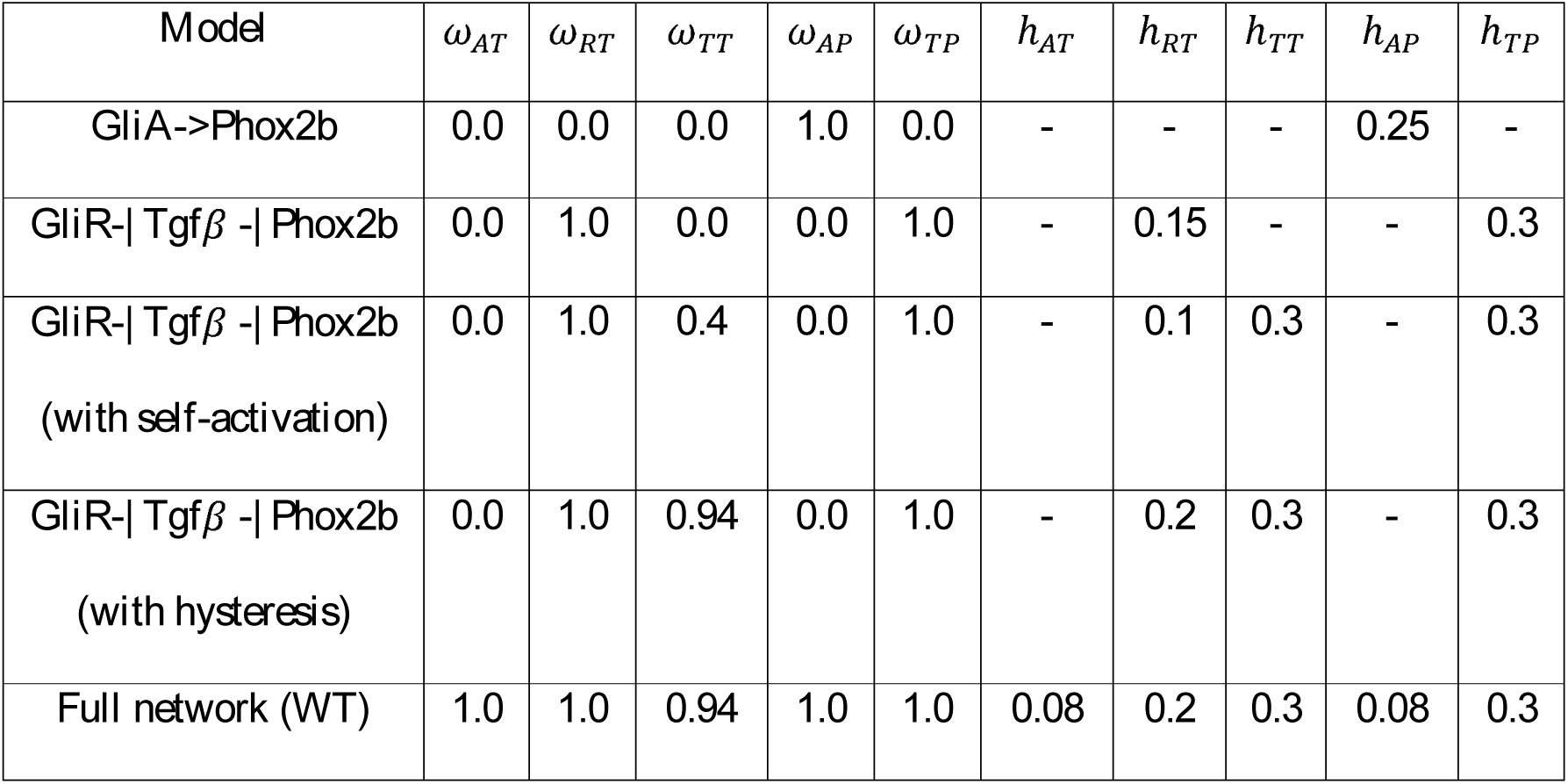

## Data availability

RNA-sequencing data has been deposited in Gene Expression Omnibus (GEO) under ID codes GSE114640 and GSE112698. All source codes are freely available at: https://git.bsse.ethz.ch/marcelob/tgfbeta_mndiff.

## Acknowledgments

We thank M. Scott, J. Eggenschwiler, K. Miyabayashi for reagents, Science for Life Laboratory, the National Genomics Infrastructure and Uppmax for assistance in sequencing and computational infrastructure.

## Author contributions

J.M.D, Z.A. and J.E. conceived the project and designed experiments. J.M.D and Z.A. performed experiments with contribution from M.K.; J.M.D, Z.A., J.E. interpreted the data with contribution from D.I.; A.J. and A.A. performed RNA-sequencing analysis. D.I., M.B. developed model; M.B. generated Fig.6; J.V. carried out preliminary modelling work. M.P.M. contributed with mutant mice. J.E., D.I., J.M.D., Z.A. and M.B. wrote the manuscript.

