## Supplementary figures for "A Shh/Gli-driven three-node timer motif controls temporal identity and fate of neural stem cells"

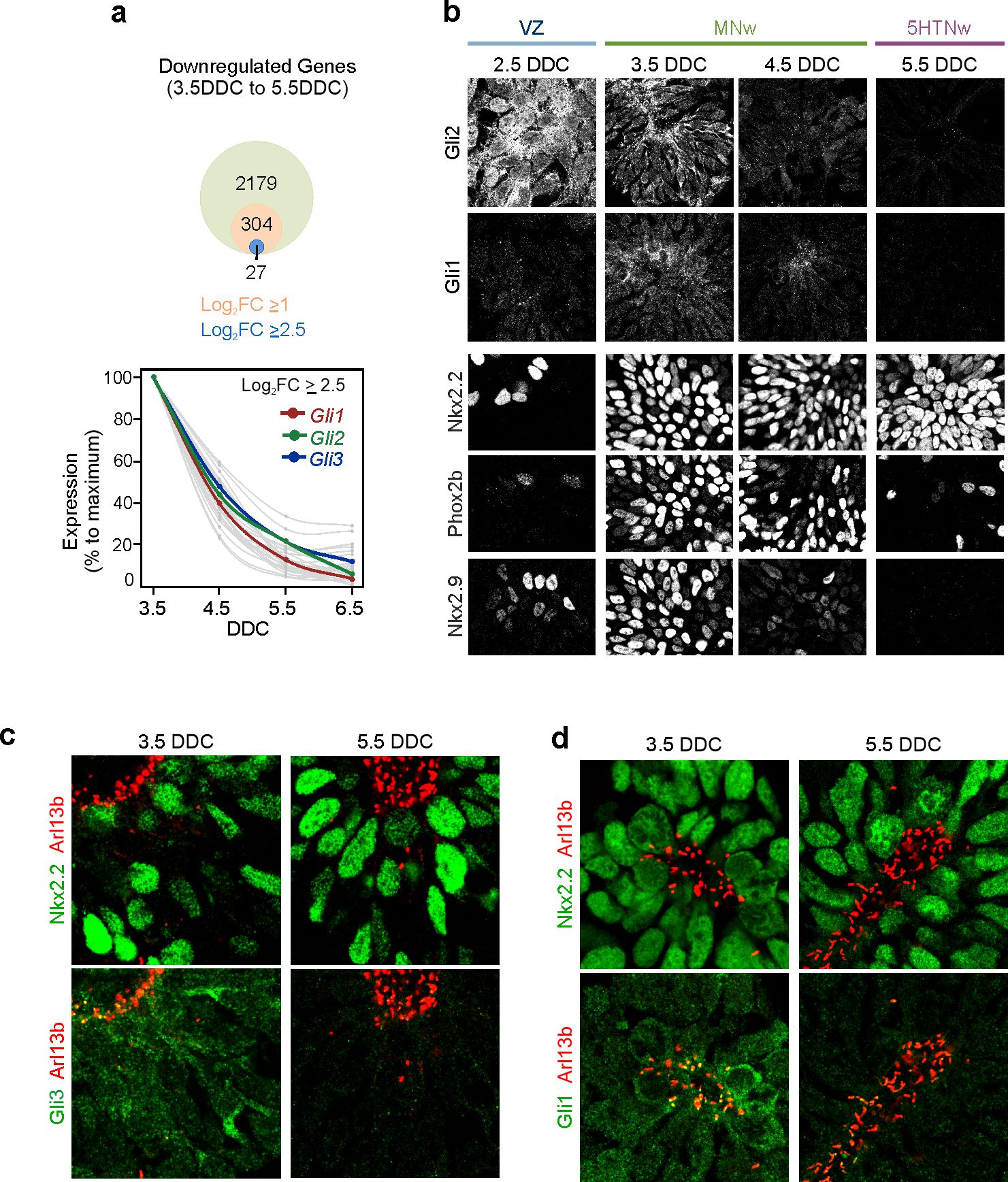

**Extended Data Fig. 1| Characterization of Gli expression in the temporal lineage.**

**a,** Genes downregulated in NSCs between 3.5 and 5.5 DCC (p≤0.05) at different fold-change (FC) cut offs, and relative temporal expression profile of genes having a log2(FC)≥ 2.5.

**b,** Immunofluorescence of Gli1, Gli2, Nkx2.2, Phox2b and Nkx2.9 in neural progenitors at 2.5-5.5DDC.

**c,d,** Immunofluorescence of the cilia marker Arl13b with Gli3 (**c**) or Gli1 (**d**) in Nkx2.2^+^ progenitors. Images presented correspond to a triple immunocytochemistry with antibodies against Arl13b and Nkx2.2 and with Gli3 (**c**) or with Gli1 (**d**), and where Arl13 immunostaining is presented with Nkx2.2, Gli3 or Gli1. DDC, days in differentiation conditions.

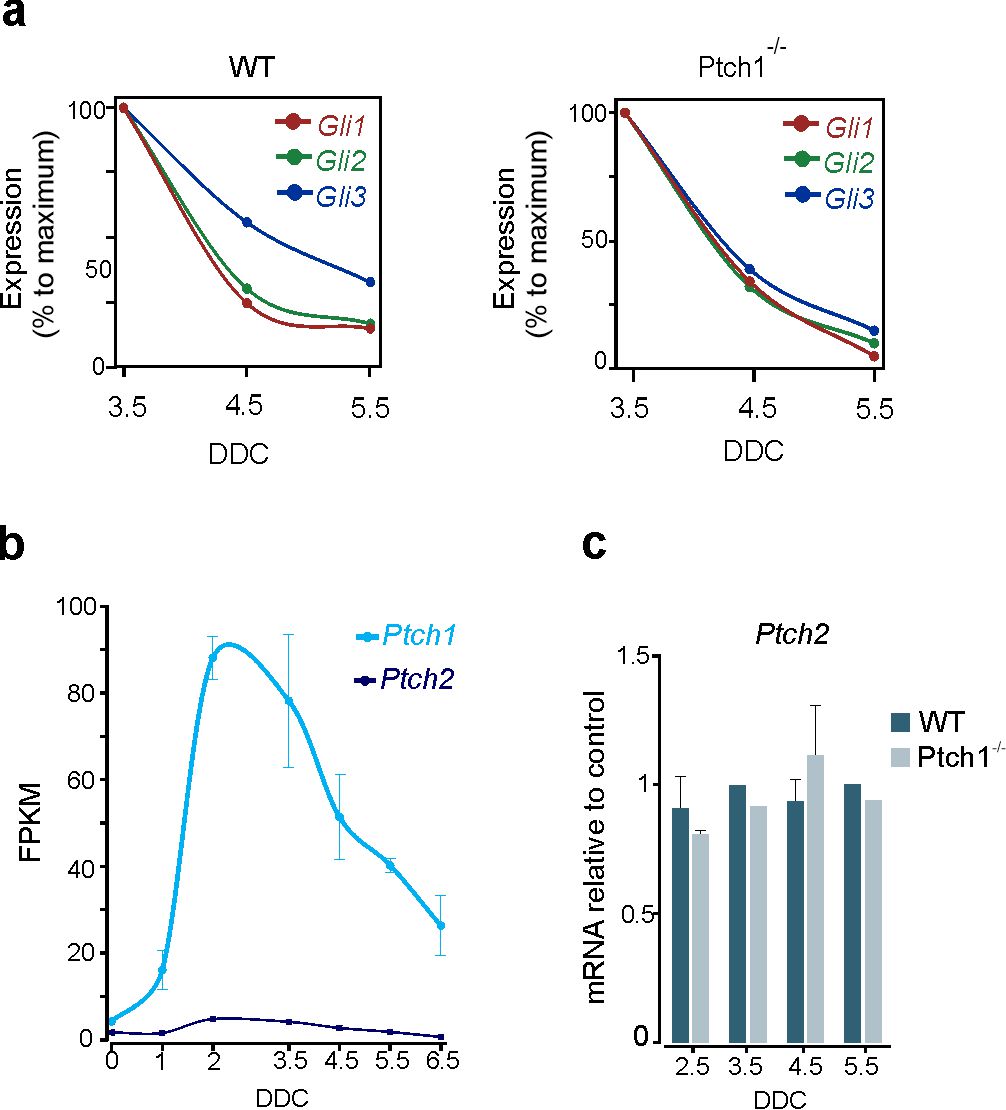

**Extended Data Fig. 2| Characterization of the expression of *Gli* and *Ptch* genes in Ptch1 mutant cells.**

**a,** qPCR analysis of *Gli1,-2, -3* gene expression in WT and Ptch1^-/-^ mutant neural progenitors isolated at 3.5, 4.5, and 5.5 DDC.

**b,** Expression levels, in FPKM, of *Ptch1* and *Ptch2* in WT neural progenitors isolated at different DDC defined by RNAseq. Values, mean ± SEM. **c,** qPCR analysis of *Ptch2* expression in WT and Ptch1^-/-^ neural progenitors during differentiation.

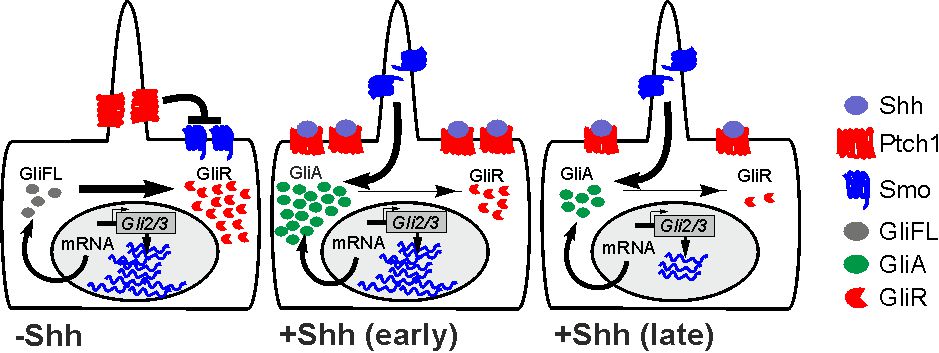

**Extended Data Fig. 3| Schematic summary of Gli dynamics in the temporal lineage**

Bifunctional Gli proteins are processed into repressors in absence of Shh. A proportion of Gli2 and Gli3 proteins are processed into GliR-forms even when Smo is fully activated, and GliA and GliR levels generated correlate to Gli transcription levels, which decrease over time.

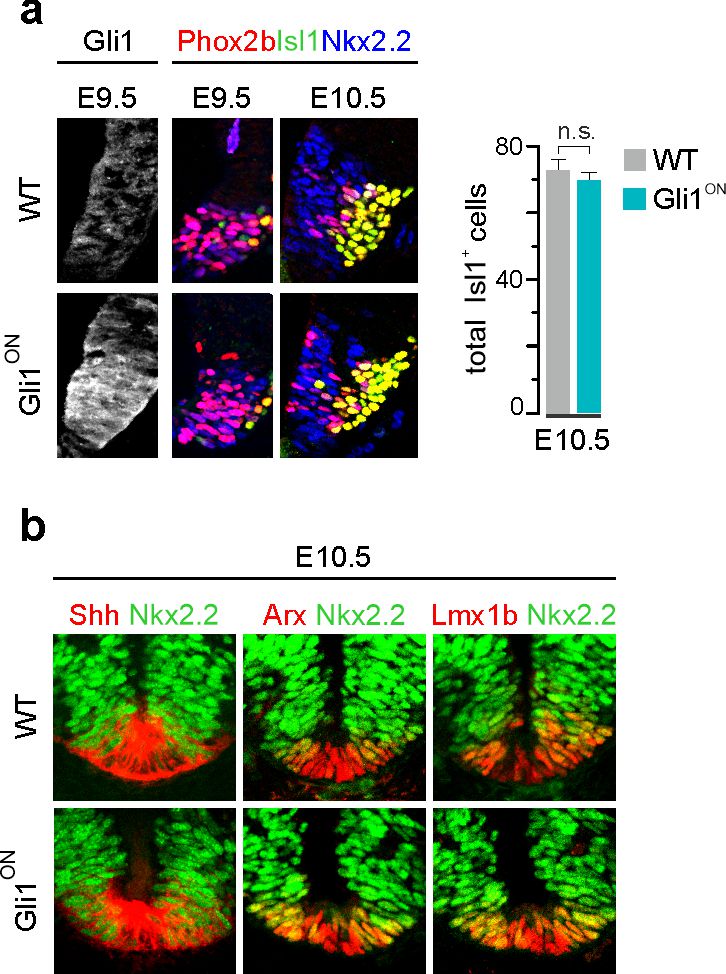

**Extended Data Fig. 4| Analysis of patterning and early MN production in Gli1^ON^ mutants.**

**a,** Immunostaining for Gli1, Phox2b, Isl1, and Nkx2.2 at rhombomere 7 of the hindbrain in WT and Gli1^ON^ embryos at E9.5 and E10.5, and quantification of Isl1^+^ MNs at E10.5. Values presented as Mean ± S.E.M.. Student´s t test, n.s. non-significant.

**b,** Expression of the FP markers Shh, Arx, Lmx1b and of Nkx2.2 at caudal levels of the hindbrain in E10.5 WT and Gli1^ON^ embryos. Nkx2.2, which is transiently expressed in ventral midline cells at early developmental stages failed to be downregulated in the FP of Gli1^ON^ mice. However, this did not affect overall FP identity

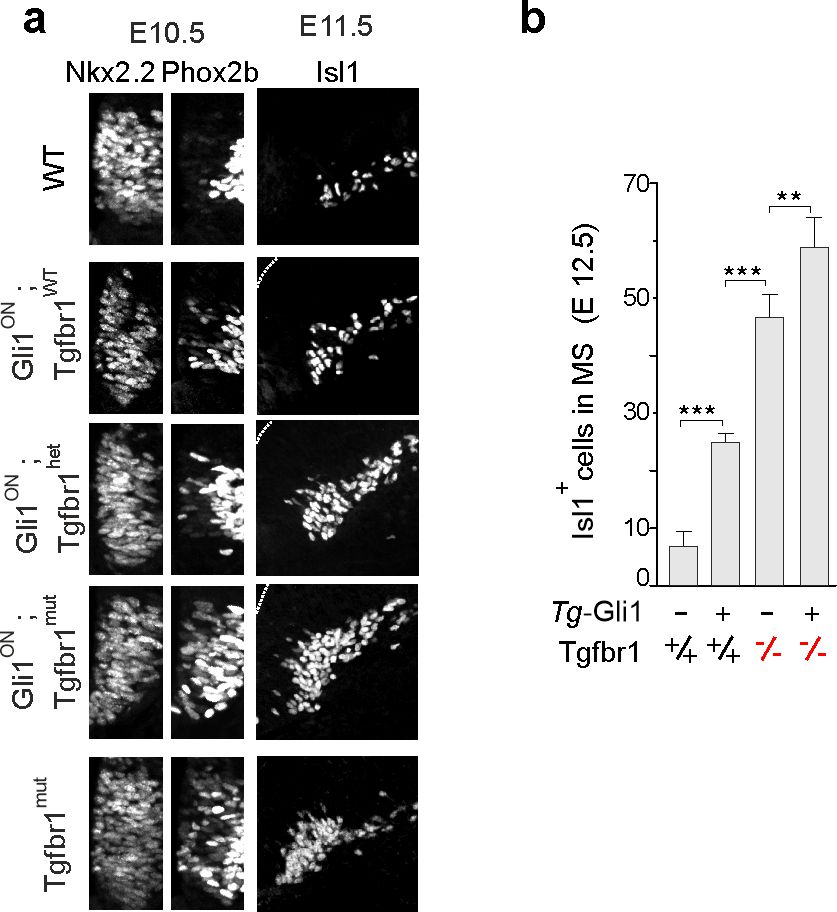

**Extended Data Fig. 5| Effects of reduction of Tgfβ signaling in Gli1^ON^ mice on Phox2b expression in progenitors and motor-neuron production.**

**a,** Transverse sections at r2/3 level of ventral hindbrain of mouse embryos at E10.5 and E11.5. Expression of Phox2b in Nkx2.2^+^ NSCs at E10.5 and Isl1^+^ motor neurons (MNs) located laterally to the Nkx2.2 progenitor domain at E11.5 in WT, Gli1^ON^, Tgfbr1^mut^ embryos and Gli1^ON^ mutants on a Tgfbr1 heterozygous (het) or homozygous (mut) background. Isl1^+^ MNs at this level migrate laterally where they settle to form the trigeminal nuclei. Thus, Isl1^+^-MN located in the trigeminal nuclei correspond to early-born MNs while those located laterally to the progenitor domain correspond to late-born MNs and can, therefore, be used as a measure of the temporal output of Nkx2.2^+^ progenitors.

**b,** Quantification of Isl1^+^ MNs in the migratory stream (MS) at E12.5 in WT (*n=*5), Gli1^ON^ (*n=*4), Tgfbr1^mut^ (*n=*5) and Gli1^ON^: Tgfbr1^mut^ (*n=*3) embryos. Error bars, mean±S.E.M.; Asterisks, Student´s t test, * p≤0.05, ** p≤0.01, *** p≤0.001.

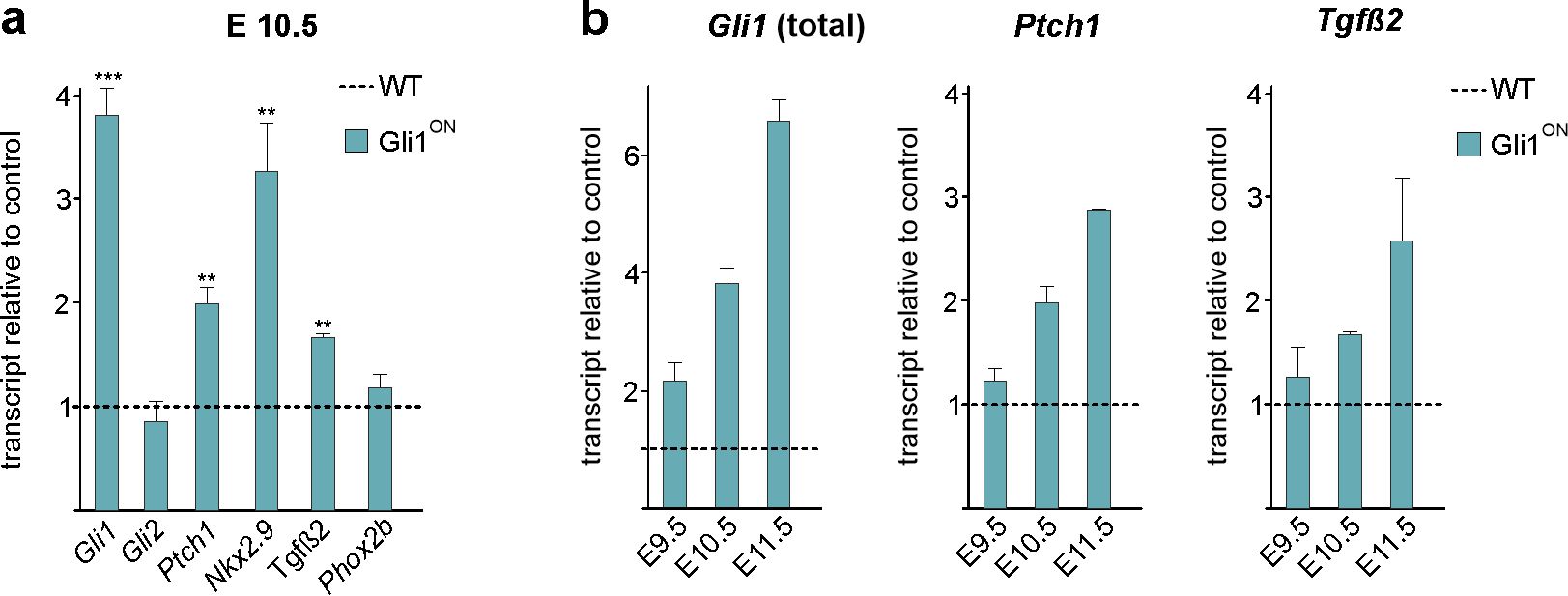

**Extended Data Fig. 6| Analysis of gene expression in Gli1^ON^ mice.**

**a,b,** Gene expression analysis by qPCR from isolated ventral neural tissue of WT and Gli1^ON^ embryos at rhombomere 7-8 level of the hindbrain. Expression of *Gli1*, *Gli2*, *Ptch1*, *Nkx2.9*, *Tgfβ2* and *Phox2b* at E10.5 (**a**) and of *Gli1* (total), *Ptch1* and *Tgfβ2* at E9.5, E10.5 and E11.5 (**b**) in Gli1^ON^ embryos relative to WT embryos. *Gli1* (total) qPCR primers detect both endogenous and transgene *Gli1*.

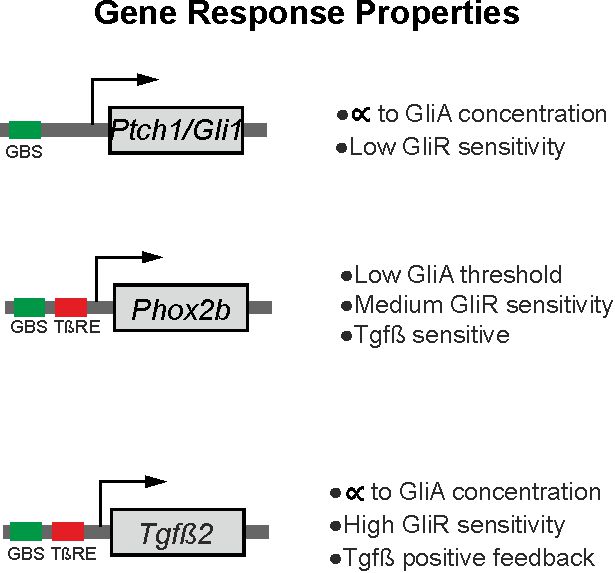

**Extended Data Fig. 7| Gene response properties**

Summary of gene response to GliA, GliR and Tgfβ signaling for *Ptch1*, *Gli1*, *Phox2b* and *Tgfβ2* genes.

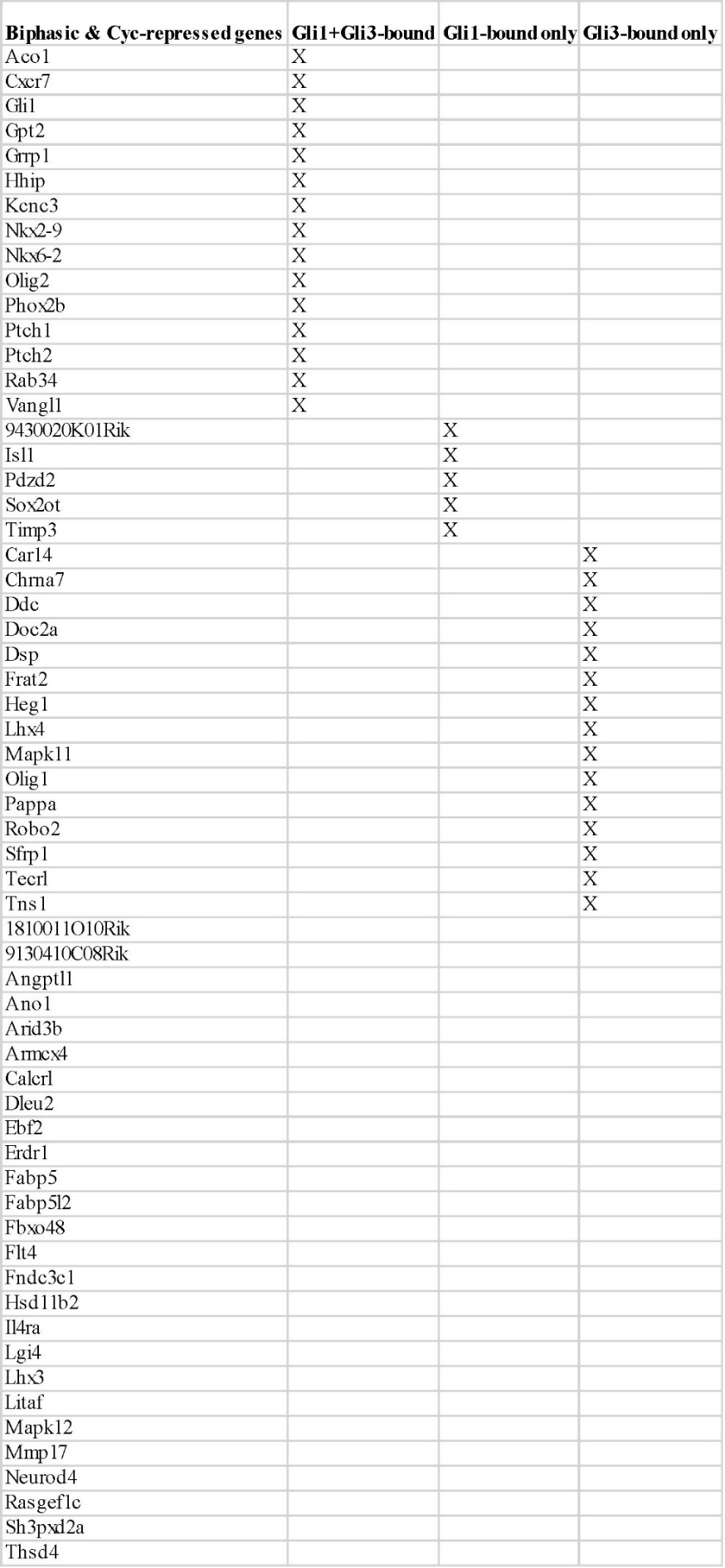
 **Extended Data Table 1|** Biphasic and Cyc-repressed genes bound by Gli1 and Gli3 proteins.
